# How neuronal morphology impacts the synchronisation state of neuronal networks

**DOI:** 10.1101/2022.12.13.520117

**Authors:** Robert P Gowers, Susanne Schreiber

**Affiliations:** Institute for Theoretical Biology, Department of Biology, Humboldt-Universität zu Berlin, Philippstraße 13, Haus 4, 10115 Berlin, Germany; Bernstein Center for Computational Neuroscience, 10115 Berlin, Germany

## Abstract

The biophysical properties of neurons not only affect how information is processed within cells, they can also impact the dynamical states of the network. Specifically, the cellular dynamics of action-potential generation have shown relevance for setting the (de)synchronisation state of the network. The dynamics of tonically spiking neurons typically fall into one of three qualitatively distinct types that arise from distinct mathematical bifurcations of voltage dynamics at the onset of spiking. Accordingly, changes in ion channel composition or even external factors, like temperature, have been demonstrated to switch network behaviour via changes in the spike onset bifurcation and hence its associated dynamical type. A thus far less addressed modulator of neuronal dynamics is cellular morphology. Based on simplified and anatomically realistic mathematical neuron models, we show here that the extent of dendritic arborisation has an influence on the neuronal dynamical spiking type and therefore on the (de)synchronisation state of the network. Specifically, larger dendritic trees prime neuronal dynamics for in-phase-synchronised or splayed-out activity in weakly coupled networks, in contrast to cells with otherwise identical properties yet smaller dendrites. Our biophysical insights hold for generic multicompartmental classes of spiking neuron models (from ball-and-stick-type to anatomically reconstructed models) and establish a direct mechanistic link between neuronal morphology and the susceptibility of neural tissue to synchronisation in health and disease.

**Significance Statement:** Cellular morphology varies widely across different cell types and brain areas. In this study, we provide a mechanistic link between neuronal morphology and the dynamics of electrical activity arising at the network level. Based on mathematical modelling, we demonstrate that modifications of the size of dendritic arbours alone suffice to switch the behaviour of otherwise identical networks from synchronised to asynchronous activity. Specifically, neurons with larger dendritic trees tend to produce more stable phase relations of spiking across neurons. Given the generality of the approach, we provide a novel, morphology-based hypothesis that explains the differential sensitivity of tissue to epilepsy in different brain areas and assigns relevance to cellular morphology in healthy network computation.

## Introduction

Network states are instrumental for neural computation: they correlate with the behavioural state in healthy animals, such as in central pattern generator circuits for movement [1, 2], and are also often altered in neuropathologies [3, 4, 5]. Recent theoretical and experimental work highlights that it is not only the connectivity between neurons which plays a role in determining network behaviour, but that neuron-intrinsic properties also exert an influence [6, 7, 2]. Mechanistically, these influences arise not only from indirect effects on connectivity, such as plastic changes in synaptic transmission or modulation of plasticity rules, but also from direct effects on the very core of processing by a single neuron: the qualitative dynamics of action-potential generation that define a neuron’s excitability class as described by Hodgkin [8]. Along these lines, weakly coupled inhibitory neurons with class 1 cell-intrinsic excitability do not foster synchronous network states [9], while neurons arranged in the same network topology with homoclinic-type action-potential generation favour in-phase synchronisation [10, 7]. Among the parameters that alter the cellular excitability class, we find those that directly affect ion channel dynamics, including channel composition, extracellular ion concentration, and modulators such as temperature [11, 12, 13, 10, 6, 14, 7]. However, these are not all.

Here, we explore the implications of a neuronal property that has received comparatively less attention in the context of network dynamics, presumably due to its more inflexible, less variable nature: neuronal morphology. Although previous work has shown that even passive dendritic arbours can play an important role in processing inputs [15, 16, 17, 18], their effect on the network state has not yet been explored. We demonstrate that differences in the extent of dendritic arborisation alone are sufficient to induce network behaviour that is either synchronised, with stable phase relationships between neuronal firing, or asynchronous. Differences in neuronal morphology thus offer an interesting and previously underestimated explanation for the differential susceptibility of neuronal tissue to synchronisation and potentially pathological phenomena such as epileptiform activity or spreading depolarisation.

Our approach exploits the fact that the dynamics of regularly spiking neurons with all-or-none action potentials come in at least three qualitatively different flavours – hereafter referred to as dynamical spiking types – corresponding to the three different mathematical bifurcations that underlie the onset of spiking at threshold. Despite the large diversity of neuronal properties, including a zoo of ion channels that shape a cell’s conductance portfolio as well as extrinsic modulators such as ionic concentrations and temperature, regular spiking in neurons is initiated by either a saddle-node on an invariant cycle (SNIC) bifurcation, a subcritical Hopf bifurcation (in conjunction with a fold of limit cycles bifurcation), or a saddle homoclinic orbit (HOM) bifurcation. The different spike onset dynamics of these dynamical spiking types have been shown to influence the temporal relationships of spikes across neurons in weakly connected networks [19, 20, 21]. Indeed, it is the combination of synaptic and cellular voltage dynamics that determines the state of the network, with the influence of cellular voltage dynamics being particularly pronounced in fast synaptic transmission. In this context, passive properties such as neuronal morphology also have an influence on the effective dynamics of spike generation. Larger dendritic branches induce a larger leak, which – as we demonstrate in this study – is sufficient to alter the dynamical spiking type, and consequently, the network synchronisation state.

Given the considerable heterogeneity between dendritic arbours — even in neurons of the same class and layer [22, 23, 24] — deciphering their influence on properties of neural computation and network states is a worthy endeavour. In this study, we methodically demonstrate that the morphology of dendritic arbours can, via their effect on a cell’s input impedance, be tuned to promote (de)synchronisation of network activity. Using a neuron model of an active conductance-based soma attached to a passive dendrite, we first recapitulate how the input impedance differs between single-compartment and dendrite-and-soma models. We then show how to calculate the local bifurcation structure (and hence the dynamical spiking type) analytically when morphology is included. Of particular interest is the Bogdanov-Takens (BT) bifurcation, which acts as an organising centre for different spiking onset bifurcations and hence dynamical types [25, 13]. In the next step, we show that the change in spike onset bifurcation from the dendritic load is reflected in the spiking timing response near the onset bifurcation. We further verify our results in detailed, anatomically reconstructed neuron models with varying degrees of dendritic arborisation, demonstrating that our results from the simplified dendrite-and-soma model are quantitatively precise, generalise and are widely applicable. Finally, we demonstrate the strong effect dendritic arborisation can have on network synchronisation via network simulations of multicompartment neurons of different dendritic extent, exemplifying the relevance of cellular morphology for the susceptibility of neuronal tissue to specific network states, including pathological dynamics.

## Results

### Passive Impedance Properties

Network behaviour is influenced by cellular properties of the constituent neurons via the dynamical spiking type they grant to each neuron. This is because distinct dynamical spiking types result in distinct timing sensitivities and consequently synchronisation properties of the network. Bringing neuronal morphology into play in a network environment therefore consists in first understanding how morphology impacts the dynamical spiking type.

Given that varying the speed at which the neuron’s membrane potential changes has been shown to induce all known dynamical switches for regular spiking [13, 10], we start our investigation by analysing how dendrites affect temporal filtering of the neuron’s membrane potential. To differentiate the voltage filtering effects caused by a dendritic arbour versus those captured by an isopotential point neuron, we first calculate the passive impedance properties of the single-compartment (S) and dendrite-and-soma models (DS). While the passive input impedance of an S model neuron is a first-order low-pass filter, spatial neuron models yield qualitatively different impedance profiles that depend on the dendritic properties [26, 27].

As we show in Figure 1, the DS model has an active soma attached to a passive dendrite which represents the equivalent cable of a branched dendritic arbour [28, 29]. If the active conductances in the soma have the same valued parameters and equations as the S model, then the two models differ only in their passive properties. The passive properties of the dendrite can be parametrised in terms of its length *L*, electrotonic length constant *λ*, passive time constant *τ*_*δ*_, and dendritic dominance factor *ρ* [29] (the ratio between the characteristic dendritic conductance and the somatic leak conductance).

**Figure 1.**
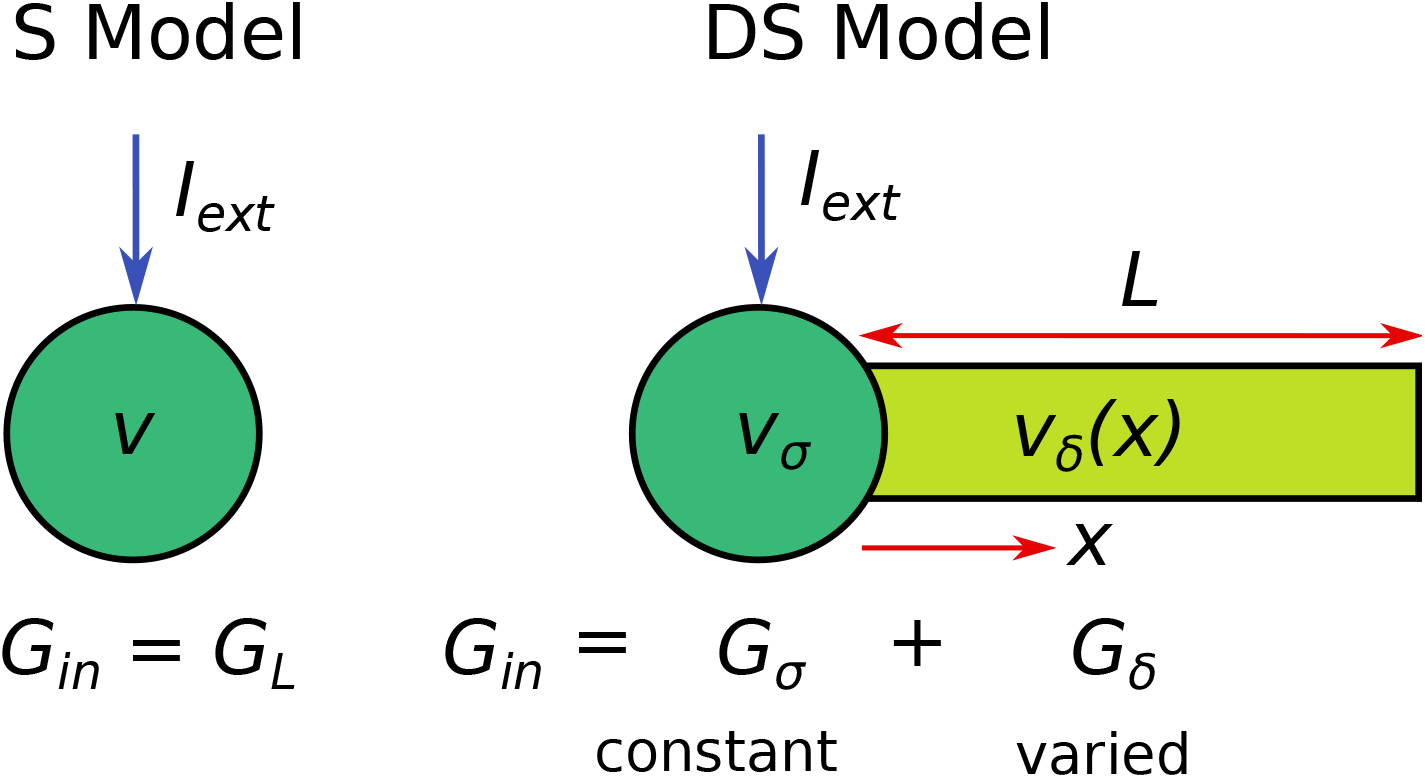
A comparison of the single-compartment (S) model with a dendrite-and-soma (DS) model with equivalent passive input conductance *G*_in_. In the DS model, the constant input current *I*_ext_ is applied to the somatic compartment. In the S model, *G*_in_ is equivalent to the lumped leak conductance *G*_*L*_, while for the DS model, *G*_in_ is the sum of the passive somatic (*G*_*σ*_) and dendritic (*G*_*δ*_) contributions. When we increase *G*_in_ in the DS model, *G*_*σ*_ is kept constant while *G*_*δ*_ is increased.

In this study, we vary the neuron’s arborisation as captured by the dendritic contribution to the passive input conductance (*G*_*δ*_). A neuron with more dendrites radiating from the soma will have a thicker equivalent cable and thus contribute more to the total passive input conductance *G*_in_. The passive input conductance is given by the constant level of external input current *I*_ext_ required to change the voltage from its steady state value (when all active conductances are zero) by an amount Δ*v*

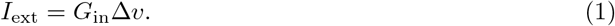

In the DS model, we apply *I*_ext_ to the soma and use the voltage change Δ*v* at the soma for *G*_in_. In each case *G*_in_ is the sum of its somatic (*G*_*σ*_) and dendritic (*G*_*δ*_) contributions. For a dendrite of effective length *ℓ* = *L*/*λ*

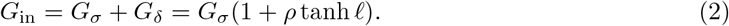

When *ℓ* ≫ 1, we can treat the dendrite as semi-infinite and the passive input conductance of the DS model becomes

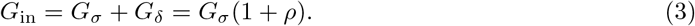

For the S model *G*_in_ = *G*_*L*_, the total leak conductance, allowing for a straightforward comparison between the two morphologies. While it is possible to adjust the dendritic parameters (*ρ, λ, L*) to match *G*_in_ for the S and DS models, this is not possible for the passive input admittance *Y*_in_ (the reciprocal of the input impedance, 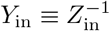). Denoting the angular frequency of the input signal as *ω*, for the S model *Y*_in_ is simply given as a first-order filter

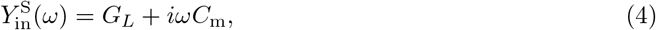

while for the semi-infinite and finite DS models we have

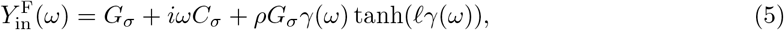

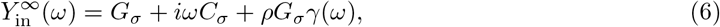

where *γ*(*ω*) is defined as

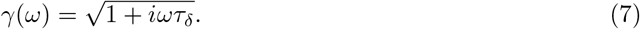

This mismatch in the passive input admittance encapsulates the difference between the S and DS models, and we will explore the implications that it has on the neuronal dynamical spiking type. For the semi-infinite DS model, *Y*_in_ can be fully parametrised by *G*_in_ and *τ*_*δ*_. From the form of *Y*_in_, we see that *τ*_*δ*_ is a key parameter for comparing the DS and S models. When *τ*_*δ*_ = 0, *γ* = 1 at all frequencies and hence 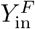 and 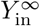 become equal to 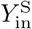 in this limit for *G*_in_ = *G*_*L*_. Thus the voltage dynamics of *v*_*σ*_ in the DS model become identical to the dynamics of *v* in the S model when *τ*_*δ*_ = 0.

We illustrate the differences between the passive input admittances of the S and DS models via its more commonly measured reciprocal, the input impedance *Z*_in_. Figure 2A shows that for any given *G*_in_, |*Z*_in_| is higher in the S model than the semi-infinite DS model at all non-zero frequencies. In Figure 2B, we see that |*Z*_in_| decreases at all non-zero frequencies when we increase *τ*_*δ*_ in the semi-infinite DS model. In the finite DS model, Figure 2C shows that if we hold *G*_in_ constant while decreasing *ℓ*, then |*Z*_in_| decreases at all non-zero frequencies.

**Figure 2.**
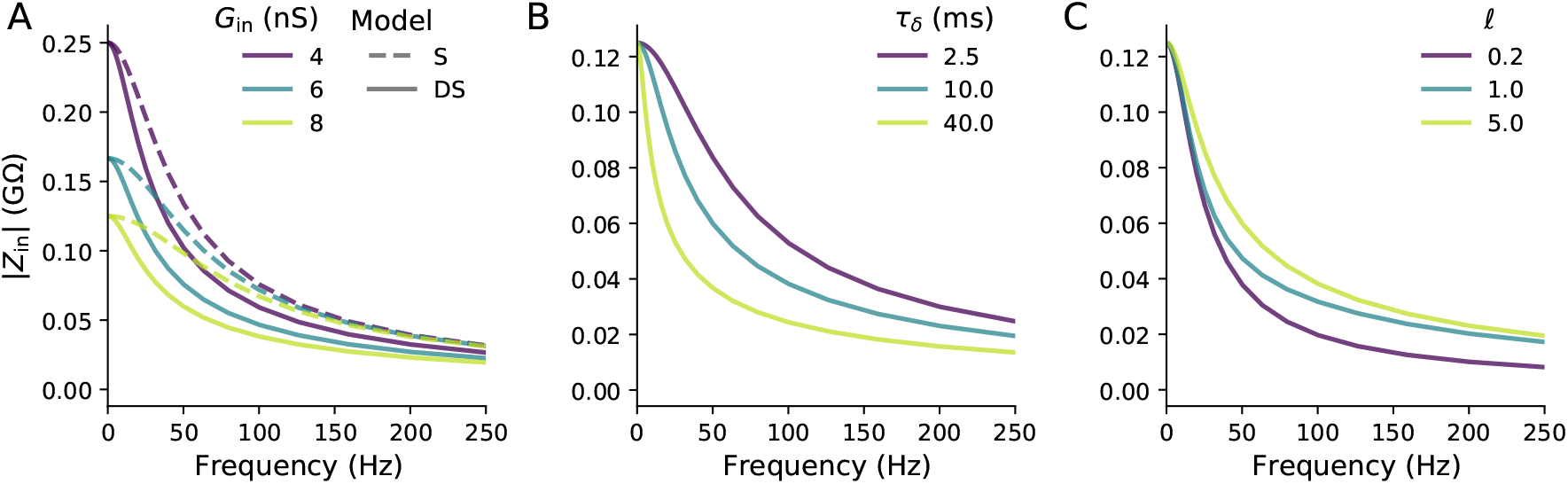
(**A**) When one fixes the input conductance *G*_in_ between the S model and a semi-infinite DS model, the magnitude of the input impedance *Z*_in_ will be higher for the S model for non-zero frequencies. *τ*_*δ*_ = 10 ms. (**B**) Increasing *τ*_*δ*_ of the DS model decreases |*Z*_in_| at all non-zero frequencies. *G*_in_ = 8 nS. (**C**) Decreasing the effective dendritic length *ℓ* while maintaining the same *G*_in_ also decreases |*Z*_in_| at all non-zero frequencies. *τ*_*δ*_ = 10 ms.

In our analyses of the DS model, we will vary directly *G*_in_ and *τ*_*δ*_ independently of each other rather than the underlying electrophysiological parameters they are defined in terms of (Equation 12). However, these parameter changes have various important biophysical interpretations. Firstly, increasing the dendritic diameter *d* increases *G*_in_ without affecting *τ*_*δ*_. Additionally, *τ*_*δ*_ can be increased without altering *G*_in_ by increasing the dendritic membrane capacitance per unit area *c*_*δ*_. Variations in the effective dendrite diameter between neurons can be significant and neuronal membrane capacitance per area may vary between mammalian species, though there are conflicting findings on this [30, 31].

While the two biophysical changes discussed so far are typically set once the neuron has developed and will not vary over short time scales, it is important to recognise that *Y*_in_ can be changed dynamically. One example of this is that increased synaptic bombardment distributed across the dendrite will increase the per-area leak conductance of the dendrite *g*_*δ*_. This in turn will increase both *G*_in_ and decrease *τ*_*δ*_ [32, 33, 34]. Increasing *d* or decreasing *g*_*δ*_ also increases the length constant *λ* (Equation 12). For short dendrites, this is an important consideration, however for long dendrites with inputs applied to the soma, *λ* does not appear in *Y*_in_ or in any of the calculations of the local bifurcations as we show in the Methods section.

### Bifurcation Structure

We now make the DS model active by giving the soma the dynamics of the Morris-Lecar model [35] while keeping the dendrite passive. The Morris-Lecar model was chosen because it has a single time-dependent gating variable, making it one of the simpler conductance-based neuron models, and also because it has been extensively studied for its ability to change dynamical spiking type upon parameter variation [19, 36, 12, 37, 13]. We chose the default class I excitability parameter set of the Morris-Lecar model with *G*_*σ*_ = 2 nS, where details of the model’s dynamics and parameters are given in the the Supporting Information.

Given that the DS model differs from the S model in terms of its passive input impedance, and that the input impedance can be fully described in terms of (*G*_in_, *τ*_*δ*_) for the semi-infinite dendrite, it naturally follows that we should choose (*G*_in_, *τ*_*δ*_) as bifurcation parameters along with *I*_ext_ in assessing the effect of morphology on the dynamical spiking type. Since the passive leak conductance in the soma *G*_*σ*_ = 2 nS, values of *G*_in_ above 2 nS denote the conductance load added to the neuron by the dendrite.

In Figure 3, we show several two-dimensional bifurcation diagrams in terms of (*I*_ext_, *G*_in_) plotted for various values of the third bifurcation parameter *τ*_*δ*_. As discussed earlier in for the analysis of the input impedance, the case of *τ*_*δ*_ = 0 is equivalent to the S model where *G*_in_ = *G*_*L*_. There are two saddle-node (SN) bifurcations at lower and higher *I*_ext_, which we will hereafter refer to as “lower” and “higher” respectively. These two SN bifurcations converge at the cusp as *G*_in_ increases. Three fixed points exist for 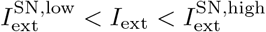, one stable and two unstable. When spiking onset is caused by a SNIC bifurcation, this occurs on the higher SN bifurcation. These bifurcations are shown in black because they do not vary with *τ*_*δ*_, and hence occur at exactly the same values of (*I*_ext_, *G*_in_) in the S and DS models.

**Figure 3.**
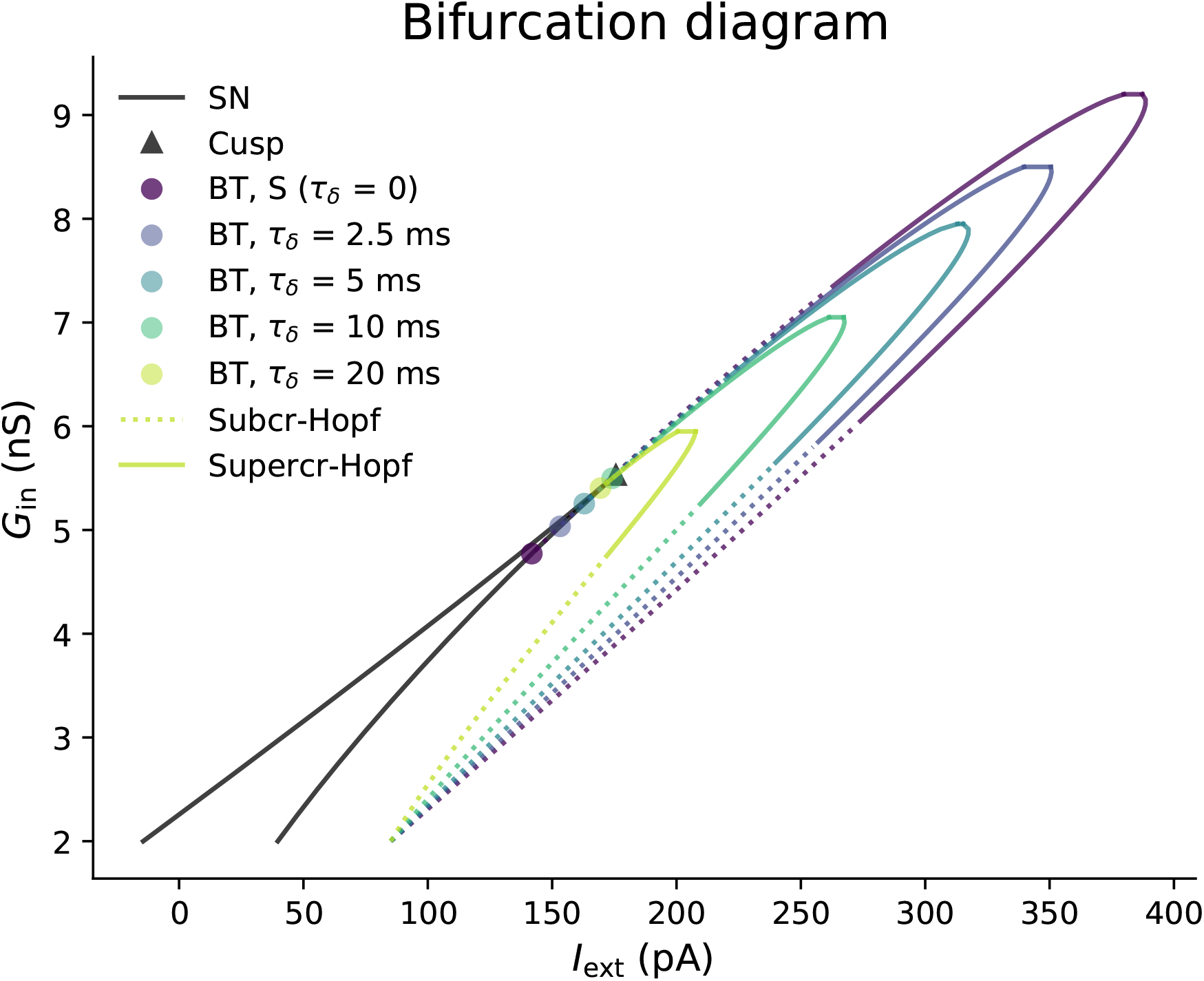
Two-parameter local bifurcation diagram in terms of (*I*_ext_, *G*_in_) of the Morris-Lecar DS model for various dendritic time constants *τ*_*δ*_. At *τ*_*δ*_ = 0, the DS model is equivalent to an S model with a leak conductance of *G*_in_. Increasing *τ*_*δ*_ shifts the BT bifurcation and shrinks the Hopf bifurcation curve. Initially increasing *τ*_*δ*_ moves the BT point to higher *G*_in_ until it reaches the cusp at the BTC point. Increasing *τ*_*δ*_ beyond 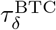 moves the BT point to the lower SN bifurcation curve and thus decreases 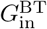. The Hopf bifurcation emerging from the BT point also switches from subcritical to supercritical when *τ*_*δ*_ passes the BTC point. The saddle-node and cusp bifurcations are shown in black because they are unaffected by changes to *τ*_*δ*_.

For *τ*_*δ*_ = 0, the BT point is located on the higher SN bifurcation and has a subcritical Hopf bifurcation emerging from it. This Hopf bifurcation permits class II excitability. Thus the BT point has often been used heuristically as separating class I and class II excitability. Increasing *τ*_*δ*_ from zero initially shifts the BT bifurcation to higher *G*_in_ until it reaches the cusp at 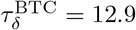 ms (hereafter denoted as the BTC point). Increasing *τ*_*δ*_ beyond 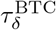 moves the BT point onto the lower SN bifurcation curve and 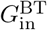 now decreases as *τ*_*δ*_ increases. As the BT point passes the cusp as *τ*_*δ*_ increases, the criticality of the emerging Hopf bifurcation switches from subcritical to supercritical at the BTC point.

When the Hopf bifurcation that emerges from the BT point is subcritical, it can be switched to supercritical by increasing *G*_in_. This criticality switch moves to lower *G*_in_ as *τ*_*δ*_ increases towards 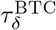. At all *τ*_*δ*_, there is a fold of Hopf bifurcations when *G*_in_ becomes sufficiently higher, and spiking is no longer possible at any applied current. The value of *G*_in_ for the fold of Hopf bifurcations decreases as *τ*_*δ*_ increases. The changes to the BT and Hopf bifurcations with *τ*_*δ*_ mean that we would expect the transition between class I and class II excitability to occur at higher *G*_in_ in the DS model for 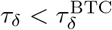.

To identify the input conductance for the SNIC to HOM switch, we must look at the saddle-node-loop (SNL) bifurcation [38, 39, 40, 10]. In Figure 4, we show the bifurcation types responsible for dynamical switching as a function (*τ*_*δ*_, *G*_in_). For all *τ*_*δ*_, 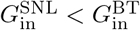, with SNIC onset for 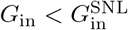. The limit cycle formed by the HOM bifurcation in this case contains all three fixed points of the system (big-HOM). Since the SNL bifurcation converges to the BTC point, it therefore makes sense that 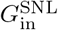 also increases with *τ*_*δ*_ when 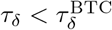. Indeed, this is what we see in Figure 4, with not only 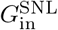 increasing with *τ*_*δ*_ but also the conductance difference between the BT and SNL points decreasing before eventually converging at 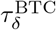. Thus both the SNIC → HOM switch and the *G*_in_ range for homoclinic onset are affected by the dendritic time constant. For 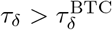, we do not find an SNL bifurcation until *τ*_*δ*_ becomes large (not shown).

**Figure 4.**
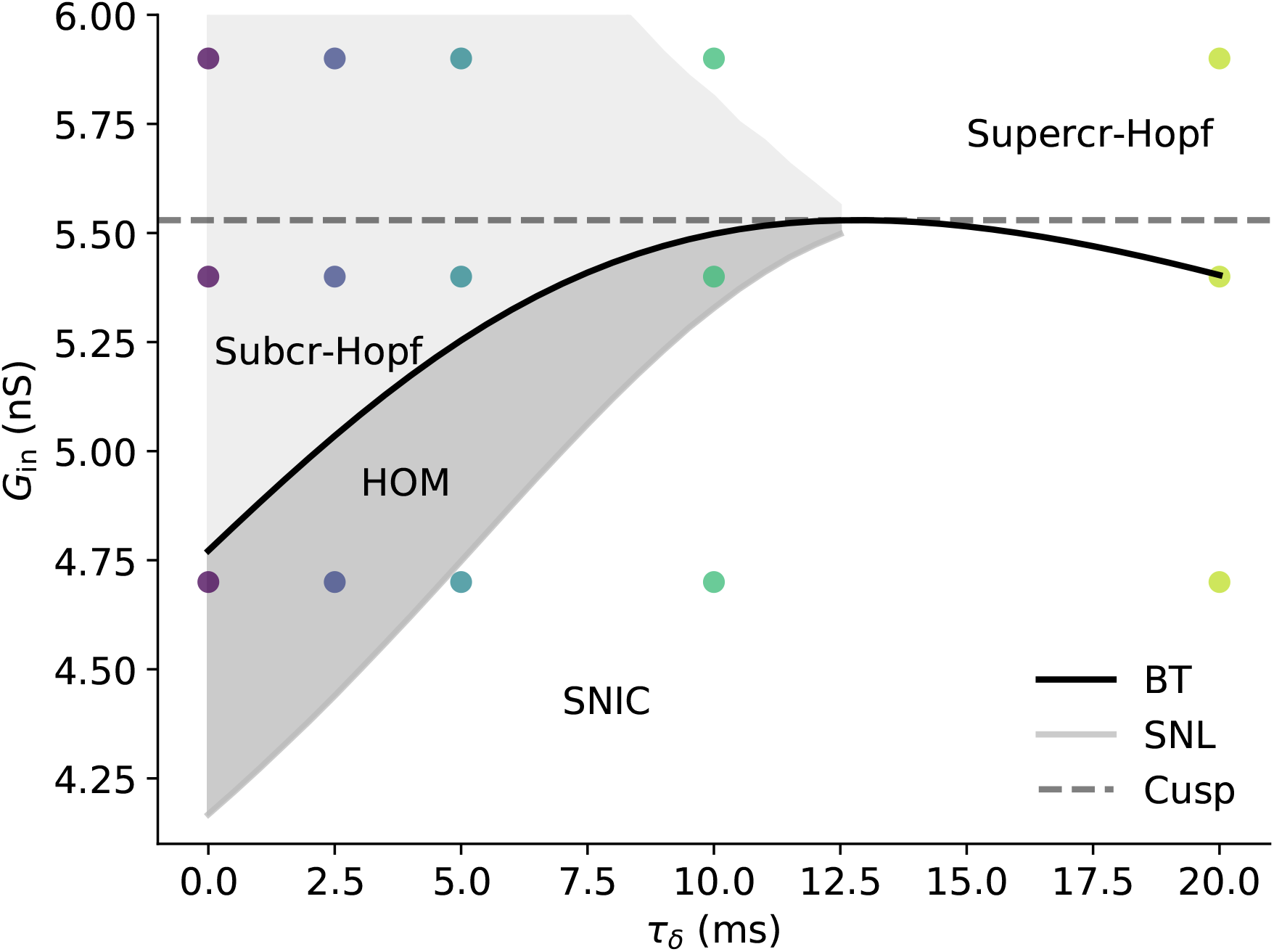
Bifurcation diagram of the Morris-Lecar DS model in terms of (*τ*_*δ*_, *G*_in_) focussing on dynamical switches. Here we have taken *I*_ext_ as the onset current for every value of (*τ*_*δ*_, *G*_in_). Points indicate values of (*τ*_*δ*_, *G*_in_) we later use for the spike timing response. At the saddle-node loop (SNL) bifurcation, the dynamical spiking type switches from SNIC to HOM, while Hopf onset becomes possible at the BT bifurcation. Increasing *τ*_*δ*_ above zero increases both 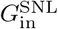 and 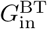 until the BT and SNL bifurcations meet the cusp at the BTC point at 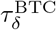. The difference between 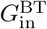 and 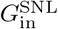 decreases as *τ*_*δ*_ increases, meaning that the range of *G*_in_ for which HOM onset exists becomes smaller. For 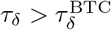, the SNL bifurcation no longer exists and 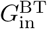 decreases. Furthermore, the Hopf onset for 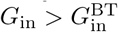 switches from subcritical to supercritical. *G*_in_ for subcritical Hopf onset increases with *τ*_*δ*_ until 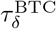, while *G*_in_ for supercritical Hopf onset decreases with *τ*_*δ*_ throughout the whole range.

Strictly speaking there can exist another global bifurcation in the region 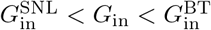 for which a fold of limit cycles (FLC) appears with a bifurcation current 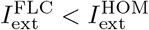, thus making the FLC the onset bifurcation [41, 13]. While this bifurcation can determine the switch between class I and class II excitability, we neglect to show it for the following reasons: (1) the spike timing perturbation response for FLC onset in this region is extremely similar to HOM onset; (2) though HOM onset in theory has class I excitability, its *f*−*I* curve is extremely steep at 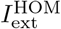, making it hard to distinguish from class II excitability; and (3) the difference between 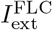 and 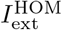 is typically extremely small [42] and thus 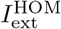 gives an accurate approximation of the onset current in this region. Thus due to simplicity and the high degree of functional similarity, we refer to the onset type for the whole of 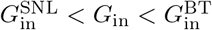 as being HOM.

We see that some bifurcations are affected by *τ*_*δ*_ while others are not. Both the SN and cusp bifurcations are calculated from the existence of fixed points alone, and so the timescale of the dynamics (such as from *τ*_*δ*_) will not affect them. On the other hand, the Hopf and BT bifurcations involve the stability switch of fixed points, for which knowledge of the timescale of the dynamics is necessary. For the SNL bifurcation, one can extend the reasoning in [10] to show that changing the timescale of the dynamics will break any existing homoclinic orbits at a given *G*_in_. As a result, one would expect that changing *τ*_*δ*_ changes the value of 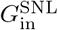.

Our bifurcation analysis indicates that switching between dynamical spiking types can be induced by increasing the dendritic arborisation as parametrised by *G*_in_. The switching between dynamical spiking types is qualitatively similar to increasing *G*_*L*_ in the single-compartment model, however the *G*_in_ values for the SNIC → HOM switch and subcritical Hopf onset increase with *τ*_*δ*_, whilst the value of *G*_in_ for supercritical Hopf onset decreases with *τ*_*δ*_.

### Phase-Response Curves

The dynamics of a neuronal spike affects how the neuron responds to external perturbations, which can come from chemical synapses, gap junction coupling, local field potentials, or externally applied currents. This has been found in both experimental [43, 44] and in modelling studies [45, 19]. The change in spike time caused by a perturbation to a tonically spiking neuron is described by the phase-response curve (PRC). In many cases, such as when neurons are weakly coupled or subject to weakly correlated inputs, one can use an individual neuron’s PRC to infer synchronisation conditions and the overall network state [19, 20, 21]. To see how the different dynamical spiking types affect the neuron’s spiking response, we calculated the PRCs for the S and DS models for a range of (*G*_in_, *τ*_*δ*_). We chose the (*G*_in_, *τ*_*δ*_) values to be in the neighbourhood of the SNL, BT, and cusp bifurcations, as shown by the coloured points in Figure 4.

At *G*_in_ = 3 nS, the Morris-Lecar neuron has SNIC spiking onset for all *τ*_*δ*_. Hence Figure 5A shows symmetric positive-valued at this input conductance. At *G*_in_ = 4.7 nS (Figure 5B), the neuron is operating in the HOM regime for lower *τ*_*δ*_ and thus we see asymmetric positively-skewed PRCs associated with HOM onset [46, 10, 47], while for higher *τ*_*δ*_ the SNL bifurcation has not yet been reached and we still have symmetric SNIC PRCs. Further increasing *G*_in_ (Figure 5C) causes the neuron with *τ*_*δ*_ = 10 ms to pass its SNL point, inducing an asymmetric HOM PRC, while the PRC for *τ*_*δ*_ = 20 ms has negative regions after passing its BT bifurcation and adopting supercritical Hopf onset. Finally when *G*_in_ is above the cusp value (Figure 5D), all the neurons have Hopf onset with negative regions in their PRCs. However, the neuron with *τ*_*δ*_ > 20 ms has a far more sinusoidal PRC with a greater negative region due to its onset being via a supercritical rather than subcritical Hopf bifurcation.

**Figure 5.**
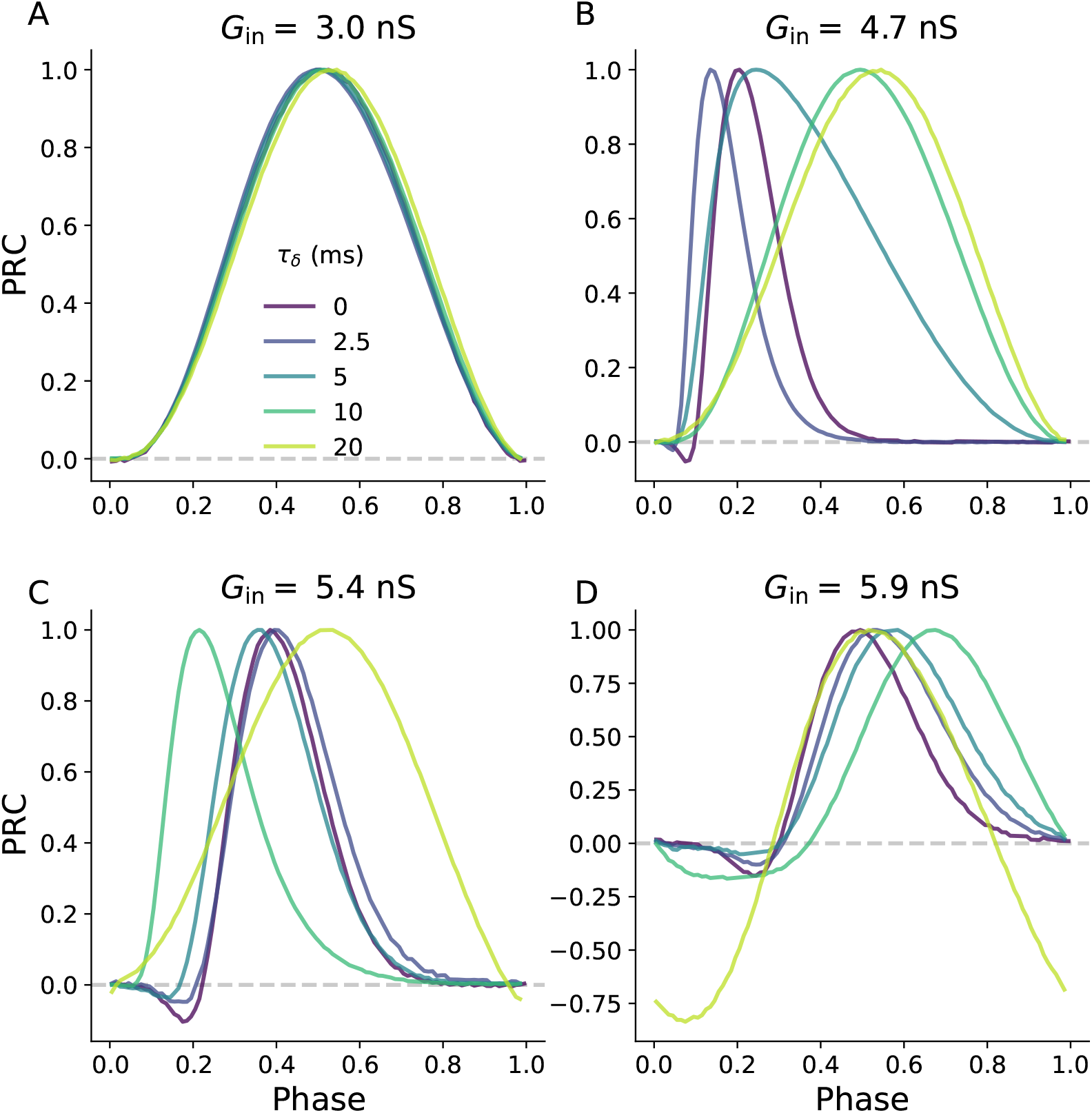
Phase-response curves (PRCs) of the Morris-Lecar DS model for various *τ*_*δ*_ and *G*_in_. Values of *G*_in_ and *τ*_*δ*_ have been chosen to be around the dynamical switches in the Morris-Lecar DS model. For example at *τ*_*δ*_ = 0, 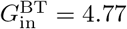 nS and for all *τ*_*δ*_ the cusp bifurcation lies at 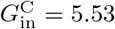 nS. When 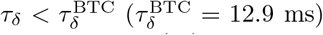, increasing *G*_in_ switches the onset PRC shape first from a symmetric SNIC PRC (**A**) to an asymmetric HOM PRC (**B-C**), and later to a Hopf-like PRC (**D**). Increasing *τ*_*δ*_ increases the value of *G*_in_ at which the SNIC → HOM transition occurs and also decreases the *G*_in_ value of the HOM → Hopf transition. For 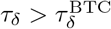, the PRC transitions straight from SNIC to Hopf-like. DS parameters used ℓ = 5.

Thus the procession of PRCs of the DS model as *G*_in_ is similar to the S model when 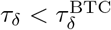. For 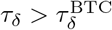, the asymmetric positive-valued PRC associated with HOM onset is not found, as predicted from the disappearance of the SNL bifurcation in the previous section. The *G*_in_ bifurcation values found earlier are thus useful predictors of the neuron’s spiking response in the dendrite-and-soma model.

### Full Morphology Test

To demonstrate that switches in the dynamical spiking type from increased input conductance are applicable to more complex and realistic neuronal morphologies, we calculated the PRCs from simulations of reconstructed dendritic arbours. In this case, we used the reconstructed dendritic arbour of a real Purkinje cell from NeuroMorpho.Org [48, 49]. We kept the somatic properties the same as the DS model investigated earlier and set all the dendritic compartments to have passive dynamics with *τ*_*δ*_ = 2.5 ms.

Starting with only the soma (representing the S model with default class I parameters), we increased *G*_in_ by adding dendritic compartments to reconstruct the full dendritic arbour in stages, as shown in the top row of Figure 6. At each stage of arborisation, we found the somatic onset current and measured the PRC. If we choose the *G*_in_ of the full dendritic arbour to be what would be in the subcritical Hopf regime of the DS model, then we can find different PRCs representing their respective dynamical spiking types by tuning the extent of dendritic arborisation. For instance, at the small arborisation in Figure 6A, *G*_in_ is low and we have a symmetric PRC indicative of SNIC onset. Increasing *G*_in_ first gives rise to an asymmetric HOM-like PRC (Figure 6B-C) before eventually yielding a PRC with substantial negative regions representing Hopf onset (Figure 6D).

**Figure 6.**
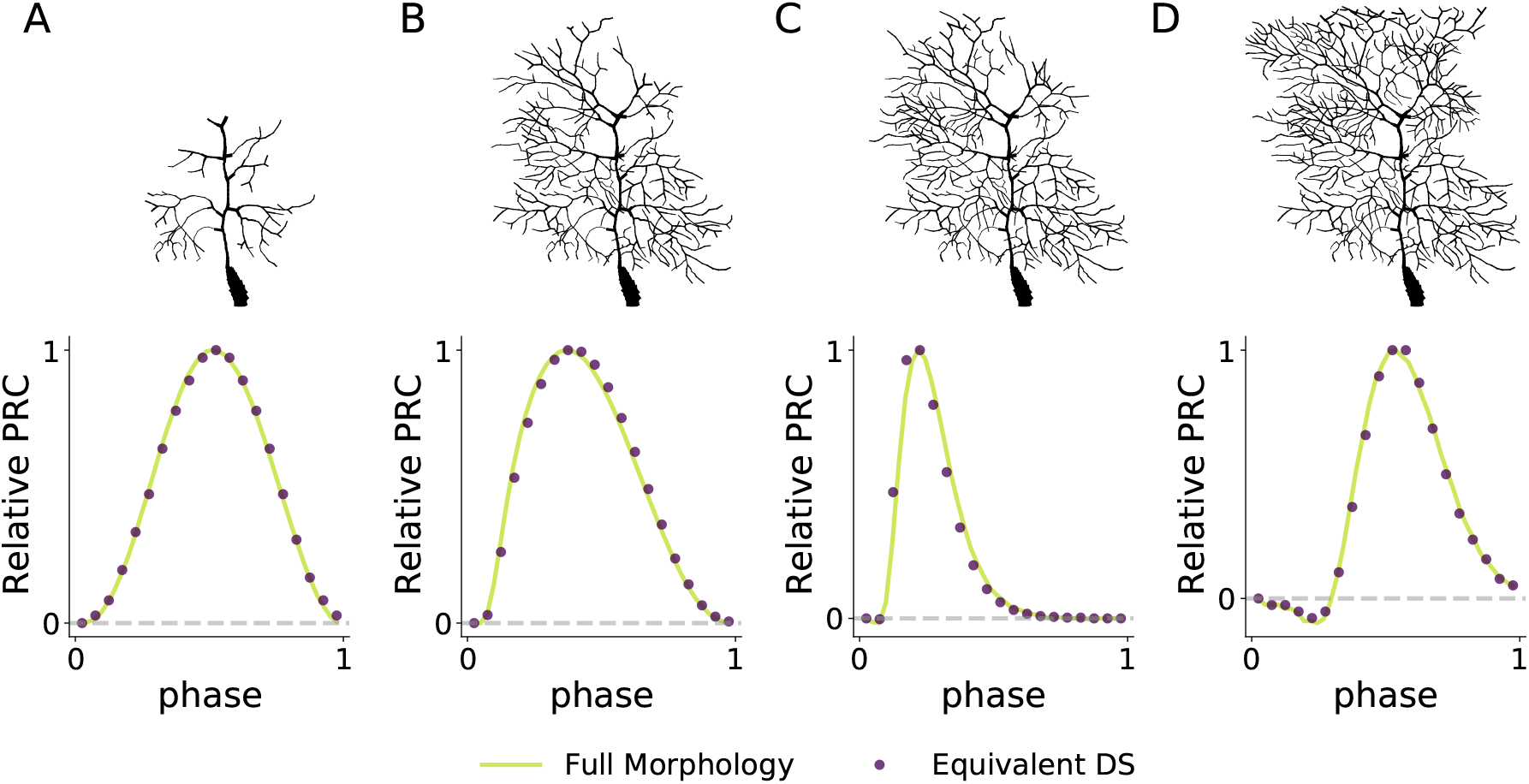
The PRCs obtained by simulating detailed multicompartment Purnkinje cell neuron show that the results of the simplified DS morphology are applicable to complicated and realistic dendritic arbours. Here we increase the input conductance of the multicompartment model by “growing” the dendritic arbour. We can see this switches the PRC shape from SNIC (**A**) to HOM (**B-C**) to Hopf (**D**) as in the DS model earlier. PRCs obtained from the equivalent DS model closely agree with the full morphology at each of the morphological stages.

We then compared the detailed-morphology PRCs with PRCs obtained from an equivalent DS model. For this DS model, the dendritic length *L* and length constant *λ* were extracted for each stage of the Purkinje cell by reducing the arbour to an equivalent cable [28, 50]. Due to the different number of compartments between the detailed arbours and their DS equivalent models, we normalised the PRCs when comparing them by setting each PRC peak value to one. Figure 6 shows that the relative PRCs of the equivalent DS model in each stage agree very closely with the PRCs obtained from the full morphology, with small deviations of PRC peak location in the HOM region being the only noticeable difference.

While we have not attempted to accurately capture the complex active dynamics of Purkinje cells (whether in the soma or in the dendrites), we have used its highly branched and complex dendritic arbour to demonstrate that our modelling of a single equivalent dendrite and soma is adequate to approximately capture the affect on neuronal dynamical spiking type of realistic dendritic arbours. Our analysis is thus not limited to the equivalent dendrite morphology in the DS model, but is applicable to realistic morphologies with highly bifurcated arbours with tapering dendrites.

### Network Simulations

We now illustrate how the dendritic conductance load can affect the network synchronisation states of a small population of these neurons via a switch in dynamical spiking type. For a group of neurons where each member has a similar spiking frequency, we measure the synchronicity of the network by the scaled standard deviation of the phase differences *ψ*_*i*_ between the *N* neurons

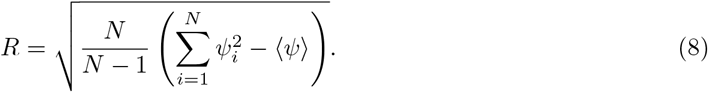

For *N* neurons all synchronised in-phase, *R* = 1, while in the splayed-out state with *ψ*_*i*_ = 1/*N* for all *i, R* = 0.

We simulated two separate networks each with 5 neurons with all-to-all connectivity of excitatory delta synapses. All synapses were connected to the soma of each neuron with zero transmission delay. In both networks the neurons had identical properties with a common dynamical spiking type: the first consisted of neurons with SNIC onset and the second of neurons with HOM onset. The synaptic strengths were chosen such that the maximum phase advance from a single synaptic input as predicted by the PRC is ∼ 0.1, as shown in Figure 7D. The firing rate of each uncoupled neuron in both networks was set at 1 Hz. Thus the two networks differ in their dynamical spiking type and not the spiking frequency or the effective synaptic strength.

**Figure 7.**
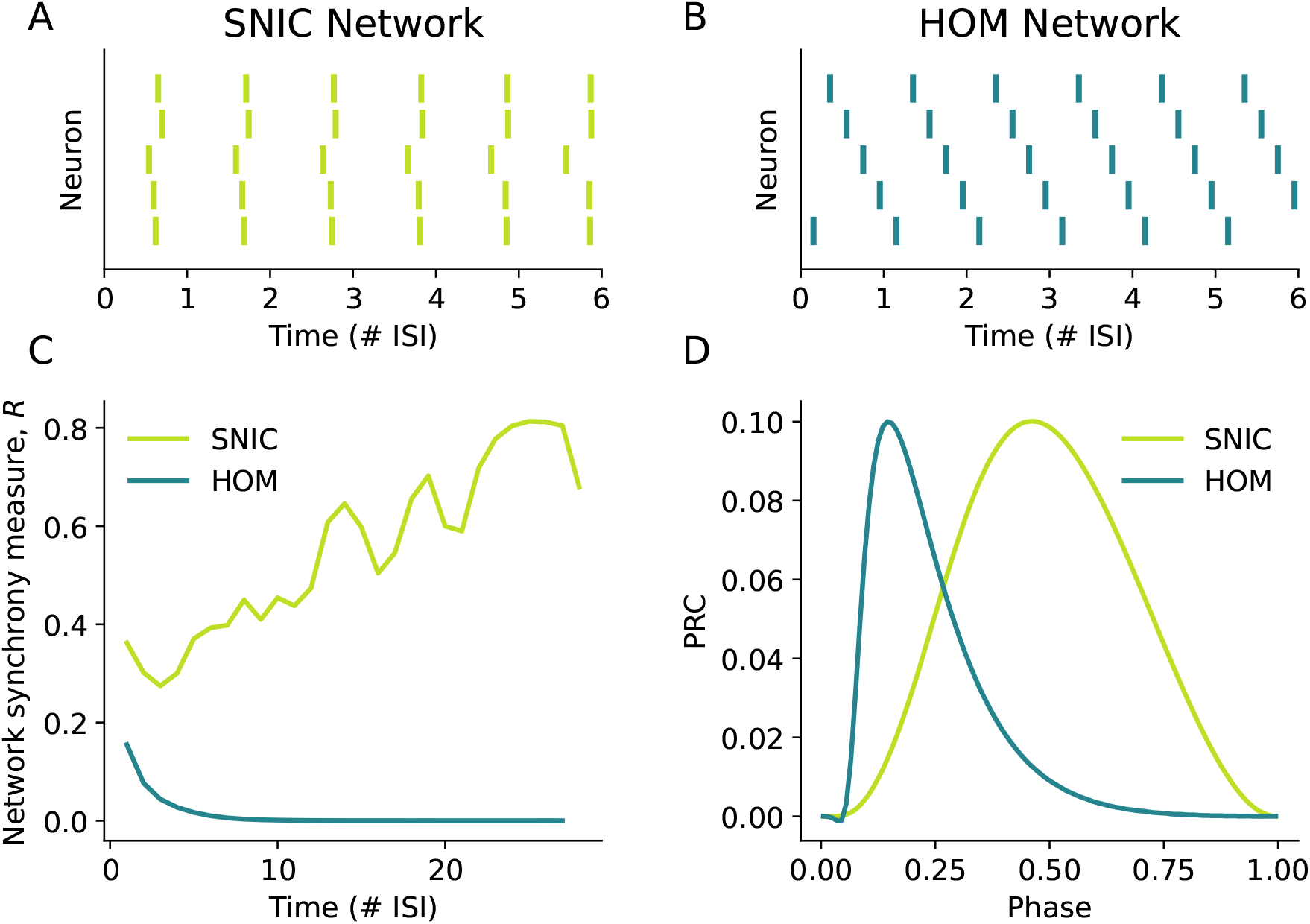
A comparison of two all-to-all networks with excitatory coupling between 5 neurons, one in which all the neurons have SNIC onset and the other in which all the neurons have HOM onset. Both networks had the same initial phase relations in the initial state with the final network states shown in (**A-B**). (**A**) The SNIC network almost achieves a synchronous network state, though this is weakly stable due to the near-symmetry of the PRC. (**B**) The HOM network achieves a splay state in which the neurons are phase-locked in which neuron *i* has a constant phase difference of 1/5 with neuron *i* + 1. (**C**) The synchrony measure over time shows that the SNIC network gradually and non-monotonically approaches the synchronous state while the HOM network converges to the splay state far more rapidly. In panels (**A**-**C**), time is measured in terms of the number of interspike-intervals (ISIs) of the network spiking rate. (**D**) The PRCs of each neuron in the SNIC and HOM networks.

The first network consisted of neurons all with low dendritic load (*G*_in_ = 3.45 nS), placing them in the region of SNIC onset. We see in Figure 7A that the phase relations between neurons approaches a synchronous state but Figure 7C shows that this network does not settle to stable in-phase synchrony. The lack of a strongly attractive stable network spiking state for this network arises from the near-perfect symmetry of the SNIC PRC [9].

By contrast, the second network consists of neurons all with higher dendritic load in the region of HOM onset (*G*_in_ = 4.53 nS). Figure 7B shows that this network converges to a splayed-out network spiking state which is approached rapidly as shown in Figure 7C. This splay state is made stable by the fact that the HOM PRC is asymmetric with a peak at a phase less than 1/*N* = 0.2 [51].

Thus we have demonstrated how changes to the neuronal dynamics conferred by a more extensive neuronal morphology can affect the network behaviour. While these network states will be altered further by synaptic dynamics, transmission delays, and heterogeneities in neuronal properties, in the weak-coupling regime the neuron’s PRC will remain an essential component in determining the stable network states.

## Discussion

In this article we have shown that not only can passive dendrites be included in the analysis of conductance-based neuron models, but also that the addition of a dendrite switches the dynamical spiking type of the neuron. In particular, we find that the reduction in input impedance caused by the dendrite can induce the SNIC → HOM dynamical switch, changing the onset PRC from symmetric to asymmetric. This dynamical spiking switch not only affects how information is processed within the individual neurons, it also implies that alterations in dendritic arborisation allow the network dynamics to achieve stable (de)synchronisation states. Specifically, our network simulations show that neurons with greater dendritic load (i.e. more extensive dendritic arbours) can achieve splay states for excitatory coupling due to their HOM onset. In contrast, these splay states could not be achieved by neurons with lower dendritic load (i.e. smaller dendritic arbours) which had SNIC onset. Different network (de)synchronisation states can thus be achieved by tuning the passive dendritic properties common to all neurons. Moreover, out findings establish a direct relationship between the susceptibility of neural tissue to synchronous network states and the morphology of the cells involved, helping us to better understand principles of neural design as well as the effect of deviations thereof in neuropathologies.

Dendritic modelling studies of conductance-based models have been performed previously. These include investigation of the effect of dendritic load on the firing frequency [52] and burst firing [15], the effect of dendritic perturbations on spike timing [53, 54], and the effect of dendritic coupling between spike-generating zones in the same neuron [55]. However, the work presented here is the first of its kind that shows via bifurcation analysis how morphology alters the dynamical spiking type and ultimately the (de)synchronisation state of the network.

The differential effect on the local bifurcations and PRCs when comparing the single-compartment and dendrite-and-soma models has utility for reducing the number of compartments in neuronal models for larger network simulations. If the input conductance is far from that of the BT point in the SNIC onset, then a single-compartment model with leak conductance equal to the input conductance of the morphology is appropriate. However, if *G*_in_ is close to 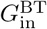, then one must use more sophisticated approximations of the input impedance (for example [56, 57, 58]). Our method showing how to calculate 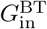 in spatially extended models thus informs one when these more sophisticated approximations are required.

Our results also prompt the hypothesis that neuronal morphology may be one crucial factor for the differential susceptibility of brain areas to pathological network states like epileptiform activity or spreading depolarisation [5, 59]. This could occur in two related ways. Firstly, pathological spiking patterns have been produced in modelling studies of single cells by changing the spiking dynamics [59, 60]. Because we have shown that the dendritic impedance alters spiking dynamics, changing the dendritic arborisation could therefore move the dynamical state of the neuron to a pathological region (via a switch in its neuronal dynamical spiking type) where network synchrony is changed. Second, pathological behaviour is often associated with either surplus or insufficient synchronous activity [3, 4, 5]. Since we have shown that the dynamical spiking switches resulting from added dendritic leak affect (de)synchronisation states, it follows that dendritic arborisation may move the degree of network synchrony to or from a pathological state.

Presently there is much research interest in how dendrites contribute to neuronal computation. This has largely focused on how either the nonlinear active dendritic channels [61, 62, 63] or nonlinear dendritic synapses in conjunction with passive dendritic compartments [64] affect the voltage or firing rate at the soma (i.e. voltage or rate coding). In demonstrating how the passive dendritic contribution changes the dynamical spiking type generated by the spiking compartment, we have added insights on one additional mechanism how dendrites can affect the temporal encoding of neuronal networks.

Methodologically, bifurcation analysis allowed us to calculate important local bifurcations of the system from a model consisting of an active soma attached to a spatially continuous passive dendrite. This approach is not restricted to the Morris-Lecar model examined in this article, but to any conductance-based neuron model with independent voltage-activated ion channels (e.g. the Wang-Buzsáki model [65]). The calculation of the BT and BTC bifurcations enabled us to predict how the dendrite affects the dynamical type, as these bifurcations act as organising centres for different dynamical types and different switches between dynamical types respectively.

Specifically, using the external input current, passive input conductance and dendritic time constant as bifurcation parameters, we found that the dendrite differentially affects the local bifurcations of the system. The saddle-node and cusp bifurcations are unaffected by the dendritic time constant of the system, meaning that all dendrite-and-soma models have the same saddle-node bifurcation locations as their corresponding single-compartment model with equivalent leak conductance. In contrast, the BT bifurcation moves in an “anticlockwise” direction about the cusp as the dendritic time constant increases, with the emerging Hopf bifurcation switching from subcritical to supercritcal at the BTC bifurcation. This change in the BT bifurcation with the dendritic time constant demonstrated that the effect of passive dendritic load on the dynamical spiking type cannot be fully replicated with a single-compartment model with an increased leak conductance.

On a mathematical note, the “anticlockwise” shift in the BT bifurcation with increasing *τ*_*δ*_ meant that the switch between class I and class II excitability occurs at higher input conductances for 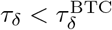. Furthermore, it means that the switch between SNIC and HOM onset at the SNL bifurcation occurs at higher input conductances, and that the input conductance region of HOM onset is smaller. Examining the PRCs confirmed the predicted spiking onset types from the bifurcation analysis: the SNIC → HOM switch is shifted to higher *G*_in_ when *τ*_*δ*_ is increased in the Morris-Lecar model. When *τ*_*δ*_ is above the BTC value, the HOM PRC region was eliminated.

Interestingly, the temporal sensitivity of neurons, as captured by the PRCs obtained from the morphologically detailed Purkinje cell reconstruction and its simplified DS model demonstrated the quantitative validity of our reduction approach in the earlier part of the analysis: PRCs of reconstructed neurons and those from the reduction of the dendritic arbour to a single equivalent cylinder were in excellent agreement. This demonstrates that the dynamical spiking type of a morphologically detailed neuron with passive dendrites can be predicted by knowing just its equivalent dendrite reduction and its active spike-generating currents. Furthermore, the equivalent dendrite reduction implies that an arbour that has more dendrites branching off the soma and thicker dendrites will have a greater impact on the neuron’s dynamical spiking type.

We note that this work also establishes a framework from which other influences of the dendritic arbour on a neuron’s spiking dynamics can be explored. As a first example, it allows us to analyse the effect of inputs applied to the dendrites. For a steady external current applied at an arbitrary location, our approach to calculating the local bifurcation structure can still be applied, as detailed in the Supporting Information. Meanwhile for transient dendritic perturbations, the PRCs from dendritic input have been simulated and observed experimentally in previous studies [17, 53, 54]. In these cases, it is found that the amplitude of the PRC decreases with the distance of the perturbation from the soma. On the other hand, the PRC shape is unchanged for positive-valued PRCs and is shifted for PRCs with negative-valued regions. Furthermore, background synaptic activity targeting the dendritic arbour also affects integrative properties of the neuron [32, 66, 67, 33, 34, 68]. This can be accommodated in our analysis via changes to the dendritic leak conductance and time constant.

The effect of some dendritic active currents can also, in principle, be included in our modelling approach. If the currents are linearisable under the quasi-active approximation [69], then the effect of the active currents can included in the filtering properties of the dendrite [70, 55, 71, 72]. Weakly active dendritic currents, such as *I*_h_ and small-conductance Ca^2+^-activated potassium current *I*_SK_, have been found to change the PRCs from being positive-valued everywhere to having negative regions [54, 73], and thus may change the network synchronisation state. If the active currents are strong enough to elicit dendritic spikes [74], then this falls outside of the framework described in our work, firstly because there are now multiple spiking compartments, and second because the interaction between strong dendritic nonlinearities (e.g. spikes, plateau potentials) and axosomatic action potentials can induce bursting behaviour outside of the regular spiking regime [75, 76, 77]. However, some aspects of our work may still be used if one considers the strongly active dendritic region to be another oscillator coupled dendritically to the axosomatic compartment [55, 78].

Lastly, there has been much interest in how the geometry of the axonal initial segment (AIS) affects spike threshold and spike shape [79, 80, 81] and encoding of time-varying input [16, 82]. Extending our framework to include features of the axon, such as the spatial separation between the soma and the spike-initiation point on the AIS, would allow exploration of how the geometry of the system affects the dynamical spiking type. Changes in the dynamical type would in turn give further insight into how axonal geometry affects the neuron’s temporal encoding of input.

In summary, we demonstrate that neuronal morphology can significantly influence the state of neuronal networks. Moreover, our results describe a method by which one can directly assess the impact of a dendritic arbour on neuronal excitability. Our approach is flexible in allowing for any conductance-based neuron model and external input currents applied to any location. Thus, the proposed approach enables further exploration of how dendritic arbour impacts the tuning of temporal encoding via changes to the network (de)synchronisation state. Our work highlights neuronal morphology as a contributor to neural function via changes to network synchrony, making it therefore likely that morphology is relevant for the differential sensitivity of neuronal tissue to synchronisation in health and disease.

## Model and Methods

### Active Soma and Passive Dendrite Model

A single-compartment (S) conductance-based neuron model consists of a leak current, *B* voltage-activated currents *I*_*a,j*_ and an externally applied current *I*_ext_

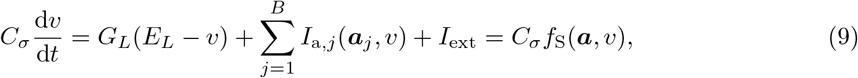

where each voltage-activated current depends on a set of activation/inactivation gating variables (hereafter termed “active variables”), 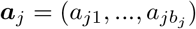. Altogether this gives *K* active variables indexed *a*_*i*_ from *i* = 1, …, *K*. As in prior research regarding conductance-based models [13, 47, 6], we assume that the active variables are independent of each other and that their steady state distributions *a*_*i*,∞_(*v*) saturate as *v* → ±∞. Each active current has a reversal potential *E*_*j*_, a maximal conductance *G*_*j*_, and depends on the product of its active variables

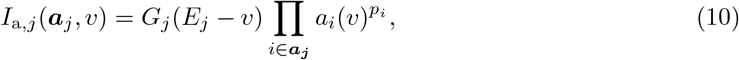

where *p*_*i*_ is the gating exponent associated with active variable *a*_*i*_. Finally, the each active variable evolves in time with a voltage-dependent time constant *τ*_*i*_(*v*)

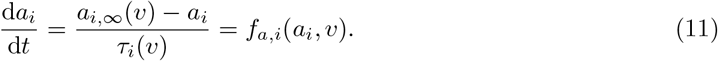

Using Rall’s principle of an equivalent cylinder, a passive dendritic arbour emanating from the soma can be simplified to a single equivalent dendrite [28, 29]. We therefore modify the above model to include the dendritic morphology by attaching a passive equivalent dendrite to a conductance-based active soma. The dendrite is spatially continuous, with the coordinate *x* denoting the distance away from the soma and *v*_*δ*_(*x*) denoting the dendritic potential at location *x*.

The dendrite is parametrised by its electrotonic length constant *λ*, its passive time constant *τ*_*δ*_ and the dendritic dominance factor *ρ* (the ratio between the characteristic dendritic conductance and the somatic leak conductance) [29]. These are each defined in terms of electrophysiological parameters

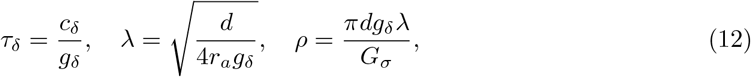

where *c*_*δ*_ is the dendritic membrane capacitance per unit area, *g*_*δ*_ is the leak membrane conductance per unit area, *r*_*a*_ is the axial resistivity, *d* is the dendritic diameter and *G*_*σ*_ is the leak conductance of the soma.

Recalling that we denoted d*v*/d*t* in (9) as *f*_S_, this means that the somatic voltage *v*_*σ*_ evolves as

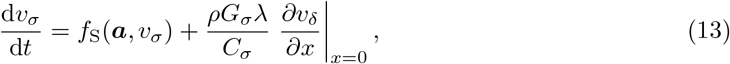

where the last term represents the axial current flowing from the dendrite to the soma.

The dendritic potential obeys the passive cable equation

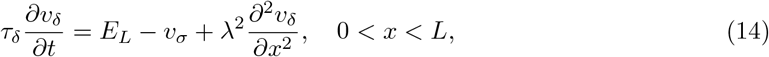

where for simplicity we have assumed that the leak reversal potential is the same in the dendrite as the soma. The dendritic potential is subject to continuity of potential at *x* = 0 so *v*_*δ*_(*x* = 0) = *v*_*σ*_, and a sealed-end boundary condition at *x* = *L*

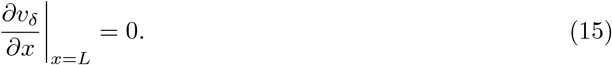

Defining the electrotonically normalised length as *ℓ* = *L*/*λ*, when *ℓ* ≫ 1, the distal dendritic end is too far away to be influenced by somatic activity, and we can simplify the model by making the dendrite semi-infinite in extent.

### Calculation of Local Bifurcations

Here we will describe how the local bifurcations of the DS system can be calculated for any conductance-based soma model with (*I*_ext_, *G*_in_, *τ*_*δ*_) as the bifurcation parameters. Although we primarily focus on the semi-infinite dendrite in this article, the method we outline here can be adapted for the finite dendrite, as given in the Supporting Information. The equations we derive for each bifurcation are applicable to a spatially continuous dendrite rather than for a specific finite number of dendritic compartments.

#### Fixed Points

Since local bifurcations are defined by the properties of a fixed point, we first outline how to calculate fixed points of the DS system. The fixed points are defined as the values (***a***\*, ***v***\*) for which all time derivatives of the system (11, 13, 14) are equal to zero. From (11), each of the active variables satisfies 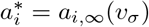 at equilibrium. The cable equation of (14) at equilibrium becomes the second-order ODE

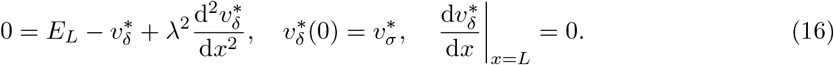

This has the following solution in terms of *v*_*σ*_

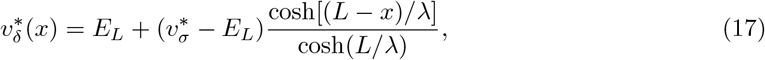

which in the semi-infinite limit reduces to

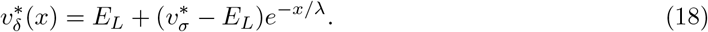

Thus all the active variables and the dendritic potential for all *x* at equilibrium are given in terms of the somatic resting potential 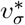. Defining the steady-state current as

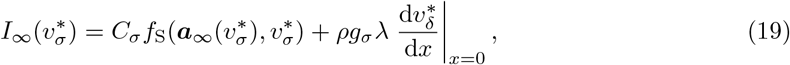

we can substitute in the dendritic voltage derivative at *x* = 0 to give in the semi-infinite case

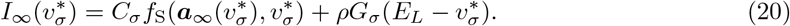

The somatic equilibrium potential 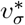 is obtained by numerically solving *I*_∞_ = 0. We note that the steady-state current equation (20) is the same as what we would find for the S model if *G*_*L*_ = *G*_*σ*_(1 + *ρ*) as *τ*_*δ*_ is not present.

### Saddle-Node (SN) Bifurcation

At a saddle-node bifurcation, a saddle and node fixed-point meet. Therefore we have a repeated root 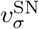 of *I*_∞_ = 0, which means at the saddle-node

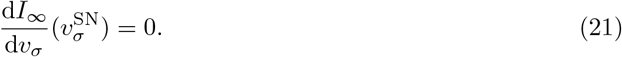

For codimension one bifurcations such as the saddle-node bifurcation, we choose *I*_ext_ as the bifurcation parameter as we are interested in the onset current for spiking. Hence we can numerically solve (21) to obtain 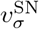 as it does not contain *I*_ext_. 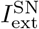 is obtained by rearranging *I*_∞_ = 0. Note that for the same *G*_in_, (21) is also identical for both S and DS models.

#### Cusp Bifurcation

At the cusp, two saddle-node bifurcations meet. Thus, the cusp bifurcation satisfies the condition

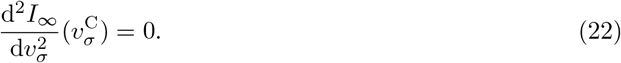

For codimension-two bifurcations such as the cusp, we will use (*I*_ext_, *G*_in_) as our bifurcation parameters. This is analogous to the prior use of (*I*_ext_, *g*_*L*_) in point-neuron models [13]. Since 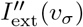 does not depend on *G*_in_ or *I*_ext_, we numerically solve (22) to obtain 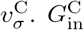 and 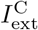 are obtained from rearranging (21) and (20) respectively. As with the saddle-node bifurcation, the cusp bifurcation condition does not depend on *τ*_*δ*_, thus for the same *G*_in_ the cusp bifurcation values will be the same for the S and DS models.

#### Hopf Bifurcation

At a Hopf bifurcation, a fixed-point changes stability and a limit cycle appears. The criticality of the Hopf bifurcation determines the stability of the limit cycle involved; at a subcritical Hopf bifurcation we have the transition: stable FP + unstable LC → unstable FP, whilst at a supercritical Hopf bifurcation we have: stable FP → unstable FP + stable LC.

The Jacobian evaluated at a Hopf bifurcation has two purely imaginary conjugate eigenvalues ±*iω*_H_. Denoting the corresponding right-eigenvector as ***q*** and its complex conjugate as 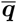, this means that we have

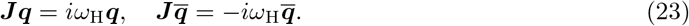

From this pair of equations, we can derive two simultaneous nonlinear equations in terms of 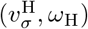 for the discretised system. Then we take the continuum limit to obtain two nonlinear equations for the continuous dendrite for which further details are found in the Supporting Information. These equations can be obtained to find 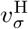 and then 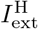 is obtained from from *I*_∞_ = 0.

Unlike all the previous bifurcations listed, the Hopf bifurcation depends on *τ*_*δ*_. Therefore, we should expect the bifurcation values of the Hopf bifurcation to differ between the S and DS models. The criticality of the Hopf bifurcation can be calculated using the approach outlined in [83], as detailed in the Supporting Information.

#### Bogdanov-Takens (BT) Bifurcation

At the BT point, a saddle-node, Hopf and saddle-node homoclinic orbit (HOM) bifurcations meet.

Since the global HOM bifurcation can also determine the spiking onset type of the neuron [40, 25, 10, 13], the BT point acts as an organising centre in two parameter dimensions for different spiking onset types. Varying the bifurcation parameters (*I*_ext_, *G*_in_) in the vicinity of the BT point can therefore induce spiking onsets associated with the three codimension one bifurcations that meet here.

The Jacobian of the system has two zero eigenvalues at the BT point. This means that ***J*** will have orthogonal left, ***l***, and right, ***r***, eigenvectors satisfying

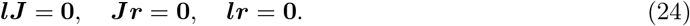

These equations can be reduced to a single equation

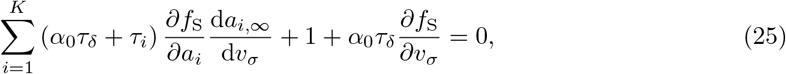

where for the semi-infinite dendrite *α*_0_ = 1/2. Since (25) has no dependence on our bifurcation parameters (*I*_ext_, *G*_in_), we can numerically solve it to find the equilibrium potential 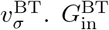 can then be obtained by rearranging the saddle-node condition (21) and 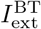 from the equilibrium condition *I*_∞_ = 0. We again see that the BT bifurcation depends on *τ*_*δ*_ and thus we should expect the location of the BT bifurcation to change with *τ*_*δ*_.

#### Bogdanov-Takens-Cusp (BTC) Bifurcation

Finally, we outline how to find the BTC point. The BTC point is of particular interest because bifurcations associated with spike onset transitions coincide at this point. At a spiking onset transition, the spiking changes from one type to another. We should distinguish this from the lower codimension BT point by stating that the BT point organises bifurcations responsible for spiking onsets, while the BTC point organises bifurcations responsible for *transitions* of spiking onsets. The bifurcations that meet at the BTC point include the Bautin (where a Hopf bifurcation changes stability), neutral saddle-node (where a homoclinic bifurcation and a fold of limit cycles meet), saddle-node loop (where saddle-node and homoclinic bifurcations meet), and BT bifurcations [83, 25, 13].

Since the BTC point acts as an organising centre for spiking onset transitions, deviations in the bifurcation parameters around it will lead to variations in the spiking onset structure in two parameter dimensions. The BTC point also has the advantage of being easier to calculate than the global bifurcations which coalesce at it, as it is calculable from the properties of fixed-points.

The BTC point has codimension 3, meaning that its location must be expressed in terms of three bifurcation parameters. In the work of Kirst et al [13], an equation for the BTC point for point-neuron conductance-based models is given with (*I*_ext_, *g*_*L*_, *C*_m_) as the bifurcation parameters, while in [6], Al-Darabsah and Campbell use the M-current conductance instead of *C*_m_. Here we use the bifurcation parameters (*I*_ext_, *ρ, τ*_*δ*_) as they are common to all conductance-based neuron models and it includes two dendritic parameters. Furthermore, *τ*_*δ*_ behaves similarly to *C*_m_, in that increasing it slows the response of the dendrite to somatic activity.

As the BTC point is where a cusp and BT bifurcation coincide, one can use the cusp condition (22) to obtain the equilibrium voltage 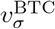, the saddle-node condition (21) to give 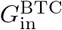, and the BT condition (25) to yield 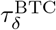.

### Simulation Details

To simulate the DS models, a cable of length *L* = 1000 μm was discretised into *M* = 50 evenly spaced spatial compartments with step size Δ*x* = 20 μm. The soma occupied a separate compartment at *x* = 0. All simulations utilised the DifferentialEquations.jl package [84] in the Julia programming language [85], from which the Tsitouras 5/4 Runge-Kutta method was used.

For the SNL bifurcation and PRCs, we required an estimate of the onset current at which stable spiking begins. Here we used the value of *I*_ext_ which produced regular spiking at the lowest possible frequency above 1 Hz. This means that for class I onset the spiking frequency was approximately 1 Hz, while for class II onset this was typically greater.

At this onset current, PRCs were obtained by increasing the somatic potential by an amount Δ*v* at a single time *t*_*k*_ corresponding to a phase *θ*_*k*_. To ensure that the phase shift is approximately in the linear regime, Δ*v* was chosen for each neuron such that the PRC maximum was never greater than 0.1. This procedure was repeated for input phases corresponding to *θ*_*k*_ = 0.01*k* − 0.005, *k* = 1, …, 100.

Code used to run all simulations, along with that used to calculate the local and global bifurcations, will be made available in the Supporting Information.

## Acknowledgments

This project has received funding from the European Research Council (ERC) under the European Union’s Horizon 2020 research and innovation program (grant agreement no 864243) and from the Einstein Foundation Berlin (EP-2021-621).

We would like to thank Jan-Hendrik Schleimer and Paul Pfeiffer for insightful discussions about excitability switching.

## Supporting Information

### Equations and Parameters of the Morris-Lecar Model

The single-compartment Morris-Lecar voltage equation is

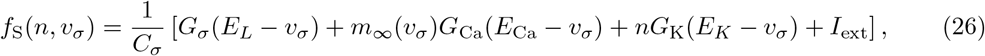

while the dynamics of the recovery variable *n* are given by

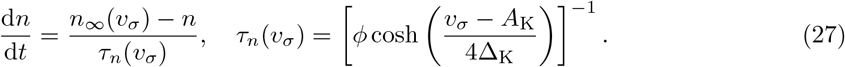

Steady-state values of the activation variables are given in terms of sigmoidal functions of *v*_*σ*_

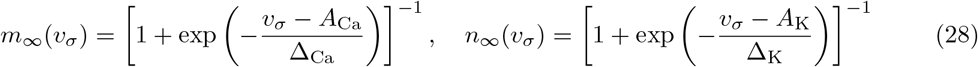

**Table 1.** Default parameters used in this paper for the Morris-Lecar model, taken from [35] and scaled by a somatic area of 100 μm^2^.

| Parameter | Value | Units |
| --- | --- | --- |
| $G_\sigma$ | 2 | nS |
| $C_\sigma$ | 20 | pF |
| $E_L$ | -60 | mV |
| $G_{Ca}$ | 4 | nS |
| $E_{Ca}$ | 120 | mV |
| $A_{Ca}$ | -1.2 | mV |
| $\Delta_{Ca}$ | 9 | mV |
| $G_K$ | 8 | nS |
| $E_K$ | -80 | mV |
| $A_K$ | 12 | mV |
| $\Delta_K$ | 8.7 | mV |
| $\phi$ | 1/15 | ms<sup>-1</sup> |

### Derivation of the Local Bifurcation Equations

#### The Method of Lines

For finite-dimensional dynamical systems, such as conductance-based point-neuron models, the Jacobian of the system has a finite number of eigenvalues. This gives a clear, but by no means trivial, approach to calculating the Hopf, BT and BTC bifurcations [10, 13, 6]. However, for a spatially continuous system, the number of dimensions is not finite. Fortunately, by using the method lines, we can derive equations for the bifurcations in an analogous fashion.

The method of lines discretises a spatially continuous system by dividing the cable into *M* compartments with index *k* = 1, .., *M* [86]. In this case, we will use a constant spatial step size Δ*x* such that *v*_*δ*_(*k*Δ*x, t*) = *v*_*k*_(*t*) and use the following discretised approximations of the first and second-order spatial derivatives

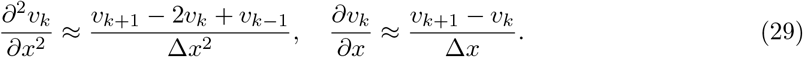

Note that *v*_*k*= 0_ = *v*_*σ*_ and that we must incorporate the sealed-end boundary condition, which changes the second derivative approximation at the distal dendritic end to

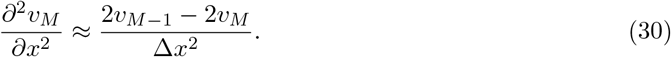

This discretisation means that we now have a *M* + *K* + 1 dimensional dynamical system, where we recall that *K* is the number of active variables. The active variable equations remain unchanged from (11), while the voltage equations in the discretised system are

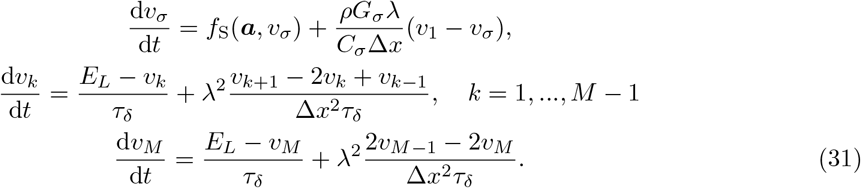

From (11) and (31) we can obtain the Jacobian ***J*** for the discretised system, allowing us to approach the problem of calculating the local bifurcations in a similar manner to the point-neuron model.

Although the Jacobian now has (*M* + *K* + 1) *×* (*M* + *K* + 1) elements, most of these are zero, and those associated with the dendritic equations do not depend on the state variables. Further details of this approach are given in the Appendix, but it is sufficient here to say that the method of lines allows local bifurcations to be calculated for the *discretised* dendrite-and-soma model. With the equations from the discretised dendrite, we can go further and take the continuum limit (Δ*x* → 0) to yield equations for the spatially *continuous* DS model. It is these spatially continuous bifurcation values which we show in the bifurcation diagrams.

With the method of lines established, we can now describe how to calculate the Hopf, BT and BTC bifurcations in a spatially continuous DS model.

#### Discretised Jacobian Matrix

Upon discretising the dendrite into *M* compartments each of width Δ*x*, we have the following *M* + *K* + 1 differential equations for the system

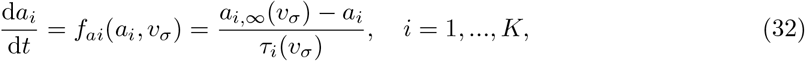

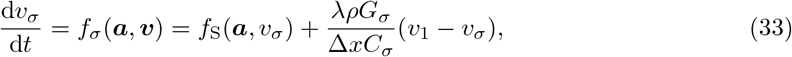

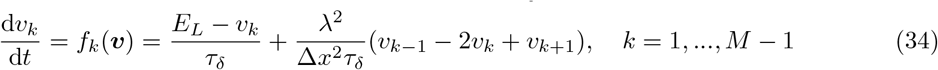

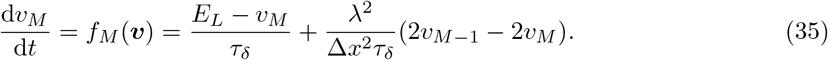

This allows us to write the Jacobian matrix as

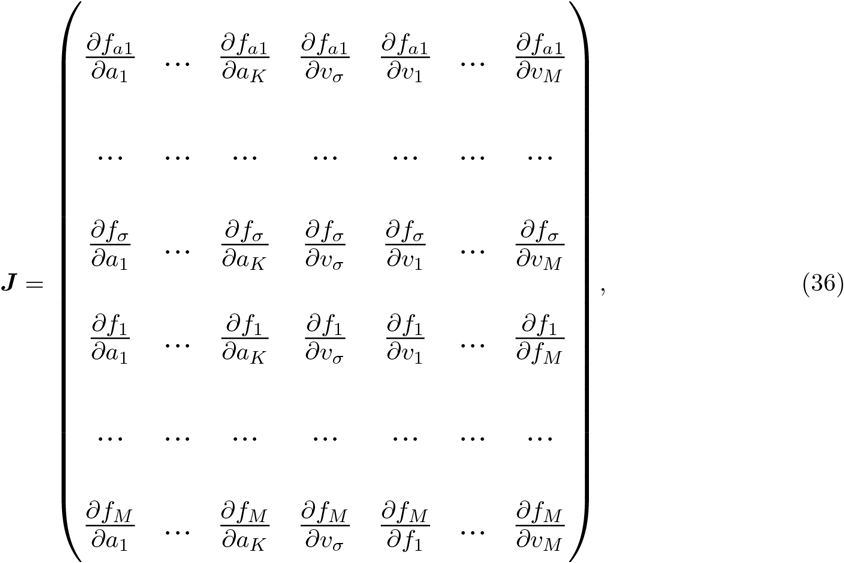

which we will split into four block matrices

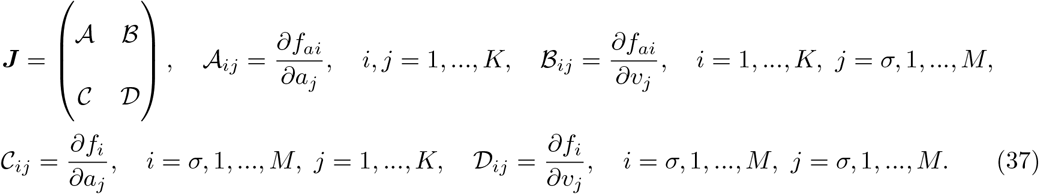

For the partial derivatives of *f*_*ai*_ we have

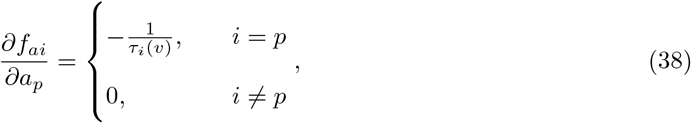

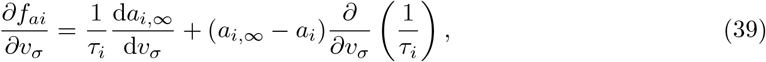

where 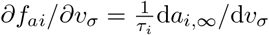 when evaluated at equilibrium. Since the active variables depend only on the somatic voltage, ∂*f*_*ai*_/∂*v*_*k*_ = 0. In terms of the block matrices, this means that 𝒜 is a diagonal matrix with

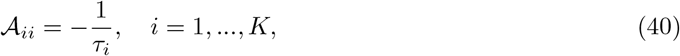

while the matrix ℬ only has one non-zero column

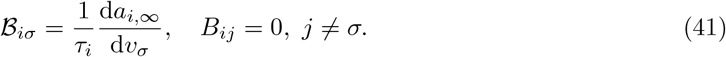

For the somatic voltage equation, we define Λ = *λ*^2^/Δ*x*^2^ and write the voltage derivatives as

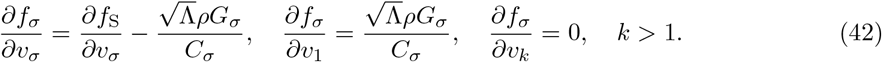

Turning to the dendritic equations, since the dendrite is passive ∂*f*_*k*_/∂*a*_*i*_ = 0 for all *i, k*. For the voltage derivatives

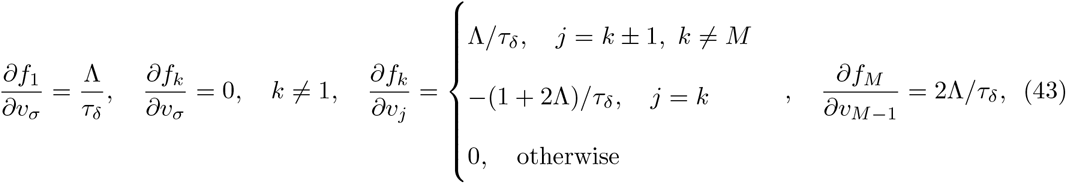

which means that the matrix 𝒞 only has one non-zero row

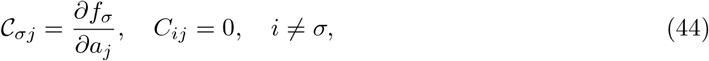

while 𝒟 is a tridiagonal matrix

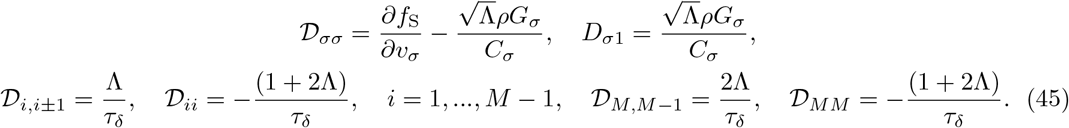

#### Bogdanov-Takens Bifurcation

At the Bogdanov-Takens bifurcation, the Jacobian has two zero eigenvalues. This means that ***J*** has left- and right-eigenvectors which not only satisfy ***lJ*** = **0** and ***Jr*** = **0** but also the orthogonality condition ***lr*** = **0**. Denoting th elements of the right-eigenvector as ***r*** = (*r*_*a*1_, …, *r*_*aK*_, *r*_*σ*_, *r*_1_, …, *r*_*M*_), the right-eigenvector equations at the BT bifurcation are

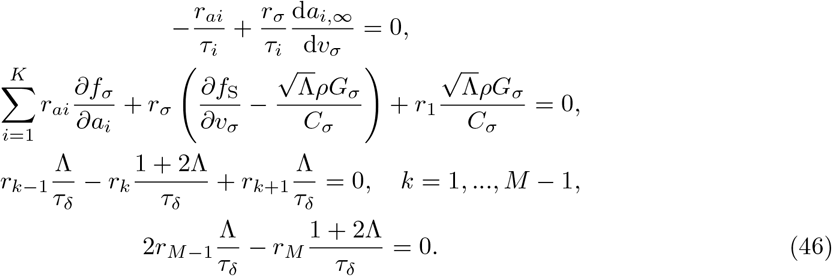

The aim here is to rewrite all the elements of the eigenvector ***r*** in terms of *r*_*σ*_, and we can immediately see that *r*_*ai*_ = *r*_*σ*_d*a*_*i*,∞_/d*v*_*σ*_. The dendritic elements meanwhile form a discrete difference equation (DDE)

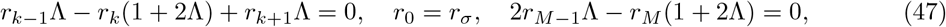

which has the general solution

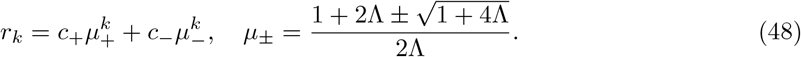

Utilising the relations *μ*_+_ + *μ*_−_ = (1 + 2Λ)/Λ and *μ*_+_*μ*_−_ = 1, we can substitute in the boundary conditions to give the specific solution as

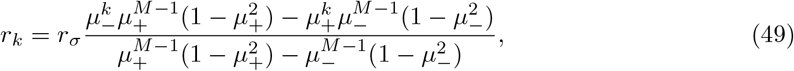

and hence we have shown that all the right-eigenvector elements can be written in terms of *r*_*σ*_. Meanwhile the left-eigenvector equations are

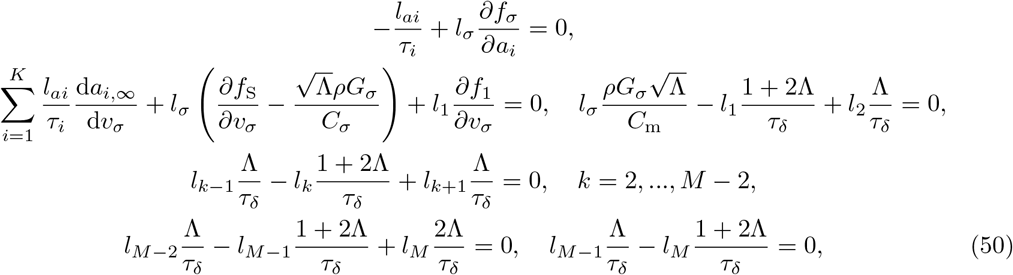

where we can see that *l*_*ai*_ = *l*_*σ*_*τ*_*i*_∂*f*_*σ*_/∂*a*_*i*_. The dendritic elements of ***l*** follow the same DDE as before, but with different boundary conditions. This has the solution in terms of *l*_1_

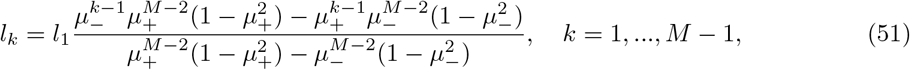

where *μ*_±_ is given by (48) and *l*_*M*_ = *l*_*M*−1_Λ/(1 + 2Λ). The dendritic elements can be written in terms of *l*_*σ*_ via

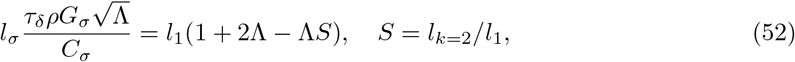

however *ρ* is intrinsically linked to *g*_in_, our bifurcation parameter. We can write *ρ* in terms of the somatic fixed-point voltage 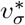 from the somatic right-eigenvector equation

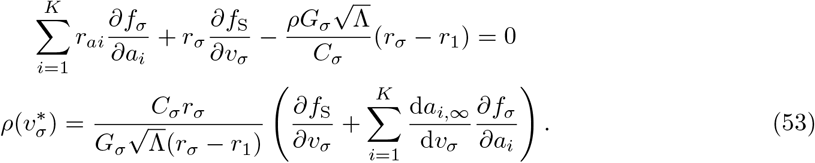

This means that each term in our eigenvector product is proportional to *l*_*σ*_*r*_*σ*_

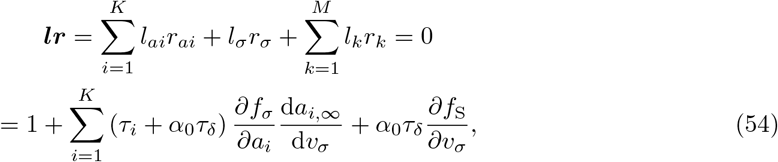

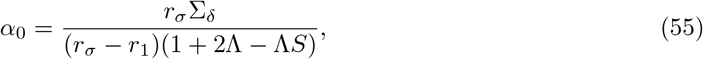

where Σ_*δ*_ represents the summation of the dendritic terms,

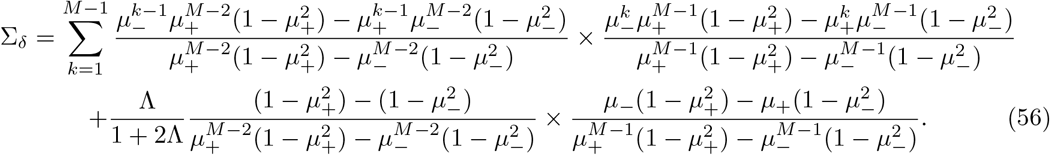

Hence for the *discrete* system, one can numerically solve the eigenvector product equation (54) to obtain 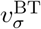 and then obtain 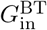 and 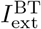 from 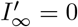 and *I*_∞_ = 0 respectively. However, we can also take the continuum limit to obtain a BT equation for a spatially *continuous* system.

A semi-infinite cable attached to a soma can be analysed by taking the limit *M* → ∞. Note that we will first consider the spatial step size to be still non-zero, Δ*x* > 0, and later we will take the continuum limit Δ*x* → 0. With an infinite number of compartments, the DDEs for *r*_*k*_ and *l*_*k*_ simply greatly as the new distal boundary conditions are

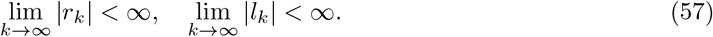

From our earlier definition of *μ*_±_, for Λ > 0 |*μ*_+_| > 1 and |*μ*_−_| < 1. This for both *l*_*k*_ and *r*_*k*_ the solutions are

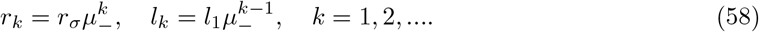

This means that the summation of dendritic terms is

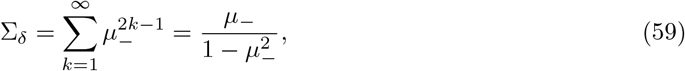

and hence the factor *α*_0_ is

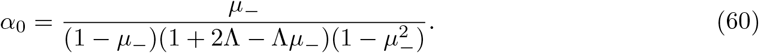

Now we take the continuum limit Δ*x* → 0 which in this case is equivalent to Λ → ∞. *α*_0_ is the only constant that depends on Λ, and has the limit

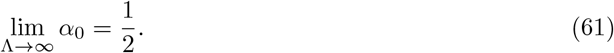

Substituting *α*_0_ into the BT condition thus gives

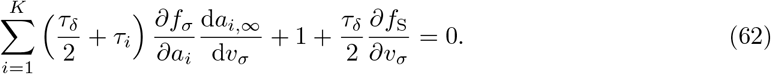

For the finite cable, after resolving the summation (56), we define Δ*z* = Δ*x*/*λ*. This allows *M* to be rewritten as *M* = *ℓ*/Δ*z* and Λ = Δ*z*^−2^. After much algebra, taking the limit Δ*z* → 0 gives for *α*_0_

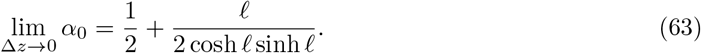

#### Hopf Bifurcation

At the Hopf bifurcation, there exist right-eigenvectors of the Jacobian ***J*** which satisfy ***Jq*** = *iω*_H_***q*** and 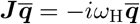. We can use the discretised Jacobian outlined earlier to obtain an expressing for the somatic voltage at the Hopf bifurcation, 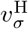. For the right eigenvectors we will use the indexing ***q*** = (*q*_*a*1_, …, *q*_*aK*_, *q*_*σ*_, *q*_1_, …, *q*_*M*_), and hence our right-eigenvector equations are

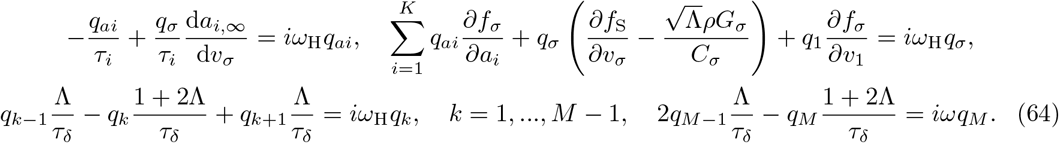

We see that the dendritic elements follow a DDE equation similar to the one for the BT bifurcation.

We rewrite the DDE and its boundary conditions as

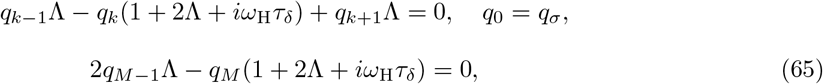

where the DDE has a general solution in terms of the bases *η*_±_, a complex analogue of *μ*_±_

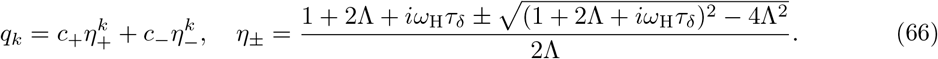

We use the boundary conditions to find *c*_±_ and hence the specific solution to *q*_*k*_

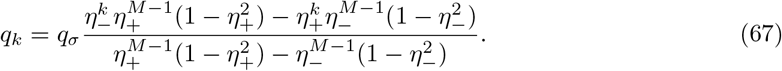

Defining 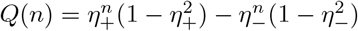, we can substitute *q*_1_ and *q*_*ai*_ into the equation for *q*_*σ*_ to yield

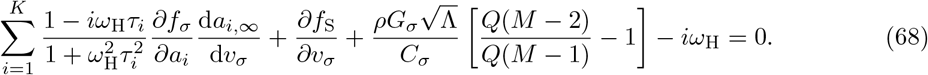

Turning back to the eigenvector 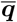, we obtain the same form of specific solution for the dendritic eigenvector elements, but this time in terms of 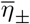

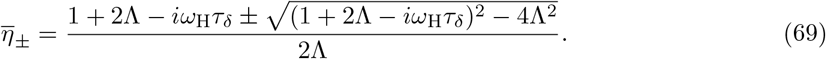

This means that the same substitutions into the equation for 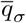 gives

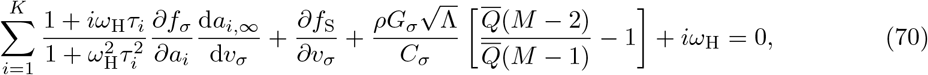

where 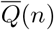 has the same definition as *Q*, but in terms of 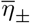.

For the semi-infinite dendrite first we take *M* → ∞, which has the effect of simplifying the specific solution of the DDEs to

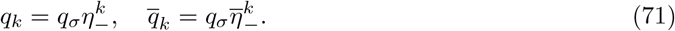

The equations for *q*_*σ*_ and 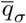 are hence simplified to

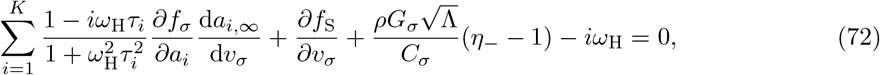

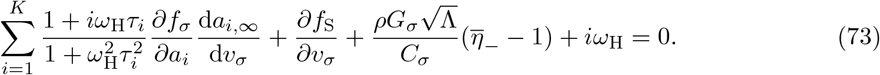

Now we take the continuum limit (Λ → ∞) of the only term that varies with Λ

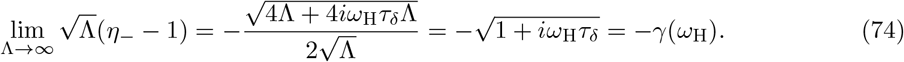

We similarly find that

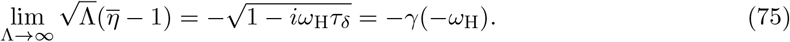

It can be shown that *γ*(*ω*_H_) and *γ*(−*ω*_H_) are complex conjugates with a sum and difference given by

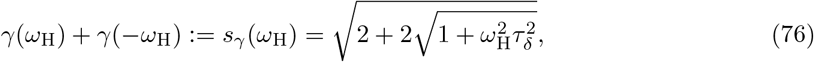

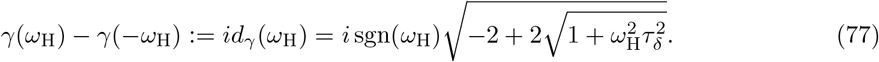

This means that summing (72) and (73) in the continuum limit gives the real-valued equation

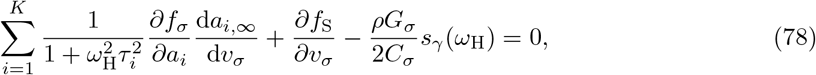

while subtracting (72) from (73) gives an equation in terms of the imaginary part

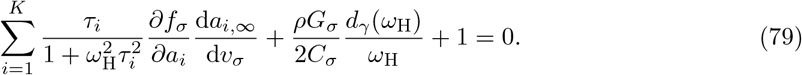

These nonlinear equations can be solved simultaneously to find *ω*_H_ and 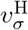.

To find the finite continuum limit solution for the Hopf bifurcation, we first find the continuum limits of 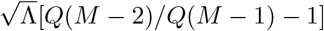 and 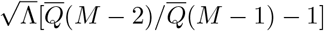 in (68) and (70) by substituting Λ = Δ*z*^−2^ and *M* = *l*/Δ*z* as in the BT section. After doing so, the first Hopf equation (68) becomes

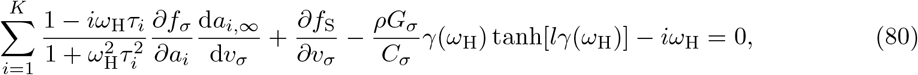

and the continuum-limit of the second Hopf equation (70) is

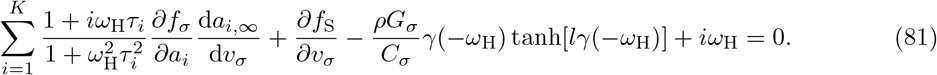

Now when we sum and difference (80) and (81) we get equations for the real and imaginary parts

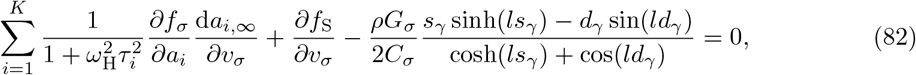

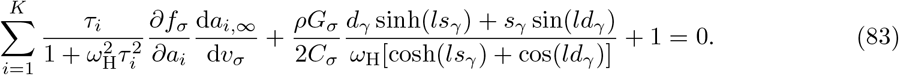

These two nonlinear equations must be solved simultaneously. In this case, we used a trust-region method implemented by the NLsolve.jl package [87].

#### Criticality of the Hopf Bifurcation

The criticality of the Hopf bifurcation can be obtained from the first Lyapunov coefficient of the equilibrium that undergoes the Hopf bifurcation

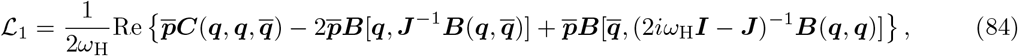

where ***q*** is the right-eigenvector of ***J*** at the Hopf bifurcation found earlier, ***p*** is the left-eigenvector of ***J***, and ***B*** and ***C*** are the second and third order tensors of the system. When ℒ_1_ < 0, the Hopf bifurcation is supercritical, while when ℒ_1_ > 0 it is subcritical. The left-eigenvectors satisfy ***pJ*** = −*iω*_H_***p*** and 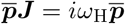 while the *i*th element of each evaluated tensor is

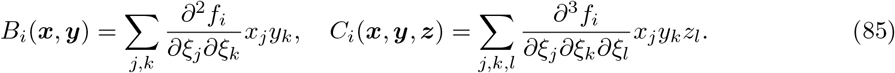

Due to the linearity of the passive dendrite, the dendritic terms of *B*_*i*_ and *C*_*i*_ will be zero. Thus, we only need calculate the *K* + 1 terms for each active value and the somatic voltage, which use the eigenvector elements *q*_*i*_, …, *q*_*K*_, *q*_*σ*_. Using the normalization 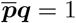, we can write all the eigenvector elements of ***p*** and ***q*** in terms of *q*_*σ*_ in the continuum limit

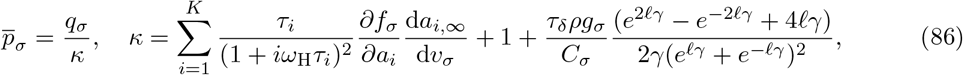

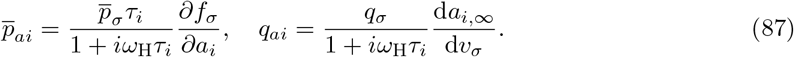

For the terms involving inverse matrices, elements required for calculation of ***B*** also have continuum limits. Denoting 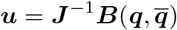, we require the elements *u*_*a*1_, …, *u*_*aK*_, *u*_*σ*_. Since only the first *K* + 1 elements of ***B*** are non-zero, this means only the calculation of (*K* + 1) *×* (*K* + 1) elements of ***J*** ^−1^ is necessary. The block structure of ***J*** defined earlier (36) means that its inverse elements can be found in terms of the blocks

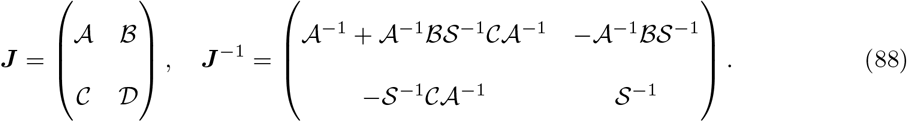

Here 𝒮 = 𝒟 − 𝒞 𝒜^−1^ℬ is the Schur complement. Using this approach, the upper-left (*K* + 1) *×* (*K* + 1) quadrant of ***J*** ^−1^ can be found as

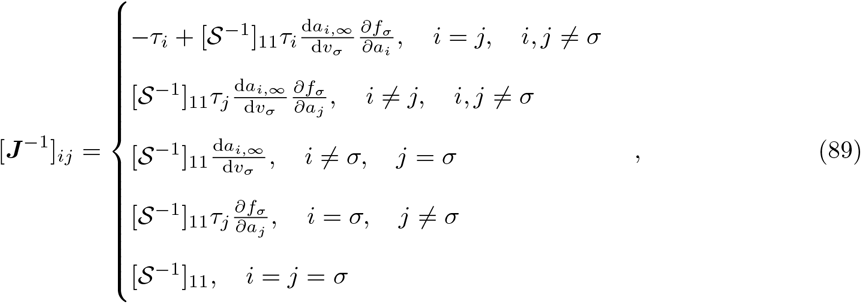

where in the continuum limit [𝒮^−1^]_11_ is the inverse of the total derivative of the somatic voltage equation with respect to *v*_*σ*_

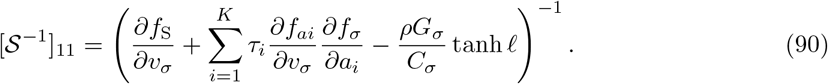

The (*K* + 1) *×* (*K* + 1) upper-left quadrant can be calculated for the inverse of 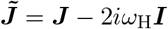 in a similar manner, yielding

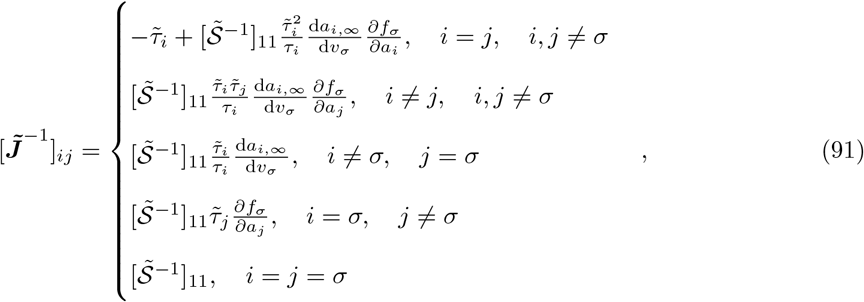

where 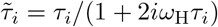 and

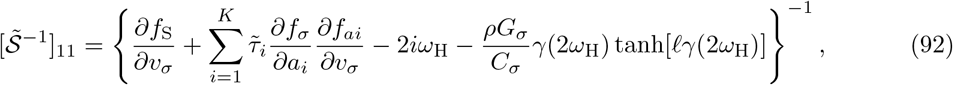

with *γ* having the same form as before. This provides us with all the information required to calculate the criticality of a Hopf bifurcation in the continuum limit.

### Dendritic Current Input

Moving the external current input to an arbitrary dendritic location *x*_in_ changes the cable equation to

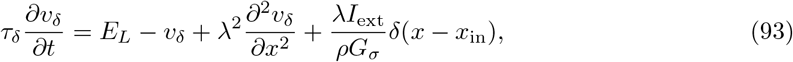

while the somatic equation remains as in (13) with *I*_ext_ = 0 at the soma. Setting all time derivatives to zero, the dendritic equilibrium voltage for the semi-infinite dendrite satisfies

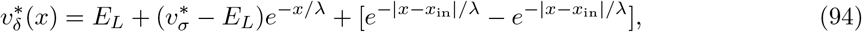

which means that the somatic fixed point potential is found by solving

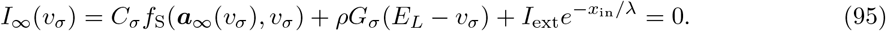

Meanwhile for finite dendrite, the dendritic equilibrium voltage with *z* = *x*/*λ* obeys

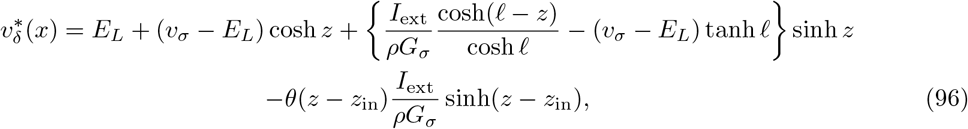

which when substituted into the somatic boundary condition yields

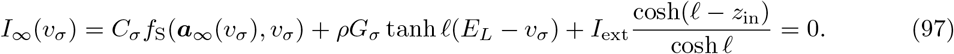

In both the semi-infinite and finite models, the only term of the steady-state current equations affected by the spatial separation between the soma at point of current injection, *x*_in_, is the external current term. The steady-state current equations converge to the somatically driven case when *x*_in_ = 0 as expected.

This means that for the saddle-node and cusp bifurcations, differentiating (95) or (97) with respect to *v*_*σ*_ removes all dependence on the location of synaptic input. Therefore, for the same values of *τ*_*δ*_ and *ρ*, 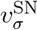 does not vary with *x*_in_ and has the same value as when external current is applied at the soma. Similarly, for the same *τ*_*δ*_, *ρ*^C^ and *v*^C^ do not depend on *x*_in_. 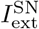 and 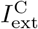 are the only parameters altered by the spatial location.

For the BT and Hopf bifurcations, we must discretise the cable equation and perform the method of lines. However, since the external dendritic drive does not depend on any of the system variables, the discretised Jacobian ***J*** will not depend *x*_in_ and will thus be identical to the somatically driven scenario. Since the bifurcation parameters for the BT and Hopf bifurcations are derived from the eigenvectors of ***J***, this means that they have values which do not vary with *x*_in_. Only *I*_ext_ varies with *x*_in_, as was the case for the saddle-node and cusp bifurcations.

### Estimation of Global Bifurcations

Global bifurcations govern many spiking onset types and spiking onset transitions, but cannot be directly calculated from the properties of fixed points. Therefore one must use numerical continuation methods [88, 89] and/or numerical simulation in order to estimate the locations of global bifurcations. For simplicity of implementation and to be understood by a wider audience, here we outline how simulation can be used to estimate the location of two global bifurcations of interest.

#### Homoclinic Bifurcation

At a homoclinic (HOM) bifurcation, a homoclinic orbit is formed at a saddle. If this orbit is stable, then it forms a spiking cycle. HOM onset is class I but its *f*−*I* curve typically grows very rapidly in comparison to SNIC onset [40]. Since the HOM bifurcation does not involve changes to the stability of fixed points, a stable homoclinic orbit typically coexists with a stable fixed point. This means that HOM onset allows bistability between quiescence and regular spiking. If the homoclinic orbit contains a single fixed point (other than the saddle), then it is termed a small-HOM (sHOM) bifurcation, while if orbit contains all three fixed points, it is a big-HOM (bHOM) bifurcation.

Using *I*_ext_ as a bifurcation parameter, if a stable HOM bifurcation exists then it often precedes a SN bifurcation, 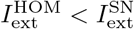. Thus we can find the HOM bifurcation as follows:

1. At a given 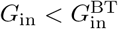, find the onset current 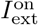 for regular spiking.
2. If 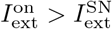 there is no bistability, and spiking onset occurs via a SNIC bifurcation.
3. If 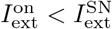, bistability exists and a HOM bifurcation is a candidate for the onset type.
4. Repeat this process at various *G*_in_ to obtain an estimate of the HOM bifurcation for the input conductance range of interest.

We specify 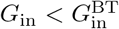 in step (1) here to avoid the presence of Hopf bifurcations. In step (3), we can only conclude that a HOM bifurcation is a candidate for the onset type because the true onset bifurcation may be a fold of limit cycles (FLC) that shortly precedes a subcritical HOM bifurcation [41, 13]. Nevertheless, this procedure forms the basis of finding the switch between SNIC and HOM onset.

#### Saddle-Node-Loop (SNL) Bifurcation

A saddle-node-loop (SNL) bifurcation is where a saddle-node and HOM bifurcation meet and has codimension two. With (*G*_in_, *I*_ext_) as bifurcation parameters, we can therefore switch between SNIC and HOM onset by varying *G*_in_ around this bifurcation. The SNL bifurcation can be estimated via the following bisection procedure:

1. Find input conductances 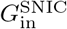 and 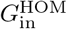 for which spiking onset occurs via SNIC and (candidate) HOM bifurcations respectively.
2. Choose a new input conductance between these two points 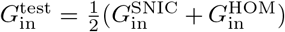 and find its onset current 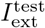.
3. Using the procedure for evaluating whether the onset is due to a HOM bifurcation, evaluate the candidate bifurcation type for onset at 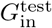.
4. Repeat step (1) with 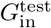 replacing the bound of its candidate bifurcation type.
5. Stop after a given number of steps or after 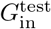 converges. The final value of 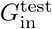 gives an estimate of 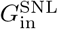.

